# Speech is defined by theta-gamma coupled acoustic rhythms, mapped onto segregated populations in human early auditory cortex

**DOI:** 10.1101/2025.10.22.683926

**Authors:** Víctor J. López-Madrona, Jérémy Giroud, Manuel Mercier, Léonardo Lancia, Bruno L. Giordano, Agnès Trébuchon, David Poeppel, Anne-Lise Giraud, Luc H. Arnal, Benjamin Morillon

**Affiliations:** Aix-Marseille Université, INSERM, INS, Institut de Neurosciences des Systèmes, Marseille, France; Institute of Language, Communication, and the Brain, Aix-Marseille Univ, Marseille, France; MRC Cognition and Brain Sciences Unit, University of Cambridge, UK; Aix-Marseille Université, CNRS, LPL, Laboratoire de Parole et Langage, Aix-en-Provence, France; Aix Marseille Université, CNRS, INT, Institut de Neurosciences de la Timone, Marseille, France; APHM, Timone Hospital, Epileptology and Cerebral Rhythmology Department, Marseille, France; New York University, New York NY, USA; Institut Pasteur, Université Paris Cité, Inserm UA06, Institut de l’Audition, Paris, France

## Abstract

Theta and gamma neural dynamics dominate the human auditory cortex during speech perception and have been proposed to track syllable boundaries and encode phonemic information, respectively. To what extent these rhythms are intrinsically generated or imposed by speech acoustics remains unsolved. Applying analytic methods from neuroscience to speech audio corpora from 17 languages, we found that canonical brain features —theta, gamma, and their phase-amplitude coupling— are a robust and specific acoustic signature of speech envelope across languages. They represent syllabic rate (2–6 Hz), vocalic features (30–50 Hz), and fundamental frequency (100–150 Hz). Intracerebral (sEEG) recordings from the auditory cortex of 18 epilepsy patients revealed that theta-gamma dynamics and their coupling are absent at rest. They emerge during speech perception and are linearly driven by the acoustic envelope, consistent with an evoked origin. Nevertheless, these responses originate from distinct yet functionally interconnected neural populations, indicating that the early auditory cortex demultiplexes speech timescales. Thus, early auditory cortex mirrors theta–gamma speech rhythms across segregated neural populations.

**IN BRIEF:** Early auditory cortex demultiplexes speech: stimulus-locked gamma activity aligns theta phase with the nested acoustic structure, functionally organising speech timescales

## INTRODUCTION

Theta and gamma oscillations in the human auditory cortex have been proposed to reflect speech encoding and parsing (Giraud & Poeppel, 2012). However, the origin and function of these rhythms remains heavily debated (Atanasova et al., 2025; Gourévitch et al., 2020; Lalor & Nidiffer, 2025). A prominent framework proposes that cross-frequency coupling — where gamma (∼25–150 Hz) amplitude is locked to the phase of theta (∼2–8 Hz; phase-amplitude coupling, PAC)— enables hierarchical parsing of continuous speech by coordinating representations of linguistic units at different time-scales (Baroni et al., 2020; Hovsepyan et al., 2020; Hyafil, Fontolan, et al., 2015; Kösem et al., 2016; Mai et al., 2016; Meyer et al., 2020; Morillon et al., 2010). However, an important ambiguity in this framework has remained unaddressed: are these neural dynamics intrinsic oscillatory mechanisms for remapping speech into internal representations, or do they mirror the rhythmic structure of the speech signal itself? The question is important because speech is not any environmental signal, but one shaped by humans, through both evolution and development, to be efficiently produced and perceived.

Research over the past two decades has extensively explored the auditory theta neural time scale (2–8 Hz) and its putative role in speech perception (Doelling et al., 2014; Luo & Poeppel, 2007; Oganian & Chang, 2019; Park et al., 2015; Peelle et al., 2013; Pefkou et al., 2017; Schmidt et al., 2023). This frequency range aligns with the primary temporal modulation in speech, approximating the syllabic rate (Coupé et al., 2019; Ding et al., 2017; Giroud et al., 2023, 2024; Greenberg et al., 2003; Kendall, 2013; Pellegrino et al., 2011; Varnet et al., 2017). Because of speech variability, it also includes suprasyllabic prominences—such as stress, accentuation, intonation contour or prosodic phrase boundaries (Tilsen and Arvanniti, 2013; Varnet et al., 2017)— usually related to the neural delta rhythm (Chalas et al., 2024; Ghitza, 2017; Keitel et al., 2017; Rimmele et al., 2021). During speech listening, the phase of neural theta activity flexibly tracks the primary speech rhythm, which is believed to aid in segmenting continuous speech into syllabic units (Ding & Simon, 2014; Giraud & Poeppel, 2012; Keitel et al., 2018; Poeppel & Assaneo, 2020). A spiking network that has a *weak* theta intrinsic activity is able to signal syllable boundaries with resilience to noise and speech rates (Hyafil, Fontolan, et al., 2015). Behavioral studies show that this tracking is crucial for comprehension (Giroud et al., 2023), which remains possible as long as the rhythm stays around the theta range (Ghitza, 2014; Lubinus et al., 2023). Thus, speech tracking within the boundaries of the theta neural scale is considered necessary, though not sufficient, for comprehension (Ahissar et al., 2001; Etard & Reichenbach, 2019; Kösem et al., 2023; Molinaro & Lizarazu, 2018; Pefkou et al., 2017).

In the auditory cortex, gamma activity has been associated with processing phonetic features and other fine-grained acoustic properties (Keshishian et al., 2023; Kösem et al., 2016; Kulasingham et al., 2020; Marchesotti et al., 2020; Mesgarani et al., 2014; Nourski et al., 2015; Teng & Poeppel, 2020). However, exploration of auditory gamma dynamics remains limited compared to theta (Baroni et al., 2020; Giraud et al., 2007; Giroud et al., 2020; Gross et al., 2013; Morillon et al., 2012). Detecting higher-frequency activity (>25 Hz) with standard surface electroencephalography (EEG) is challenging due to low signal-to-noise ratio (SNR) and limited spatial sensitivity, often requiring MEG or intracranial EEG for more accurate measurements (Baillet, 2017; Mercier et al., 2022). Additionally, low-gamma (∼25–50 Hz) and high-gamma (>50 Hz; i.e., high-frequency activity; (Leszczyński et al., 2020; Mukamel et al., 2005; Quyen et al., 2010) signals are often mixed up. High-gamma reflects broadband activity linked to multi-unit firing, whereas low-gamma exhibits narrowband oscillatory properties (Buzsáki et al., 2012).

Current models propose that low-gamma activity in the auditory cortex enables fine-grained sampling of spectro-temporal details required for phonetic processing (Ghitza, 2011; Giraud & Poeppel, 2012). This “fixed sampling” hypothesis posits that phonetic features are best encoded through an endogenous oscillatory process operating at a stable rate (∼25–50 Hz, in the low-gamma range), independently of the input dynamics. This mechanism is thought to work in tandem with theta-based syllabic parsing, via PAC, enabling efficient dual time-scale speech processing (Ghitza, 2011; Giraud & Poeppel, 2012; Hyafil, Fontolan, et al., 2015). In this framework, theta cycles adapt to the speech rhythm, while low-gamma activity represents a relatively steady sampling process optimized for extracting phonetic features within the syllabic structure (Doelling et al., 2014; Ghitza, 2013; Hyafil, Fontolan, et al., 2015).

While computational models support this dual-time scale framework (Baroni et al., 2020; Hovsepyan et al., 2020; Hyafil, Fontolan, et al., 2015), experimental evidence remains limited. There are mixed findings of intrinsic low-gamma activity in the (left) auditory cortex, with a few studies reporting it (Giraud et al., 2007; Morillon et al., 2010), while others do not (Groppe et al., 2013; Hillebrand et al., 2016; Keitel & Gross, 2016; López-Madrona, Trébuchon, Bénar, et al., 2024). Furthermore, no experimental study has clearly identified which acoustic or linguistic features engage these neural auditory oscillations, nor has established how they are encoded through PAC.

Here we aim to experimentally address the origin and function of theta and gamma activity in speech processing by characterizing the quasi-rhythmic properties of speech acoustics and early auditory cortical activity, and assessing the functional relationship between them. Similar to the theta time scale —first identified as a neural rhythm during natural speech processing (Luo & Poeppel, 2007) and later recognized as mirroring the primary acoustic speech rhythm itself (Doelling et al., 2014; Howard & Poeppel, 2012; Oganian et al., 2023; Y. Wang et al., 2025)— we hypothesize that low- and high-gamma activity may also correspond to acoustic properties of speech, driving neural responses at these frequencies. Specifically, we ask: (i) Do theta and gamma activity in the auditory cortex correspond to *intrinsic* neural rhythms, or are they driven by the structure of the speech signal? (ii) Does theta-gamma coupling (PAC) in the auditory cortex also mirror the phase and frequency of speech acoustics? and (iii) If so, is this coupling a neural mechanism for hierarchical segmentation of speech?

We analyzed speech corpora from 17 languages as well as intracerebral (stereotaxic EEG, SEEG) recordings from the auditory cortex of 18 epilepsy patients, both at rest and while they listened to a naturalistic French narrative. Our findings reveal that (i) theta and gamma dynamics and their interaction (PAC) are inherent features of the speech envelope; (ii) PAC is a robust and specific acoustic signature of speech envelope across languages; (iii) these acoustic rhythms reflect syllabic rate (2–6 Hz), vowel features (30–50 Hz), and fundamental frequency (100–150 Hz); (iv) alpha-like (6–9 Hz) oscillations dominate the human auditory cortex at rest, whereas theta and gamma dynamics are absent; (v) theta and gamma responses during speech perception are directly and linearly driven by speech acoustics; (vi) these responses arise from distinct neural populations within overlapping regions, reflecting functional segregation; and (vii) directed connectivity reveals a gamma-to-theta influence, challenging the classical theta-to-gamma framework.

Thus, the early auditory cortex appears to act as a temporal demultiplexer, preserving the hierarchical temporal structure of speech for downstream linguistic processing.

## RESULTS

### Theta-gamma phase-amplitude coupling (PAC) is a distinctive acoustic signature of speech across languages

We first performed a corpus analysis of 17 languages spanning multiple typologically different language families (mean duration per language = 5.85 hours; Table S1; see Methods). Across languages, the power spectrum of the speech envelope showed a prominent peak around 5 Hz (between 4 and 6.5 Hz across languages; Figure 1a), consistent with previous reports (Chang et al., 2025; Ding et al., 2017; Giroud et al., 2023, 2024; Varnet et al., 2017). At higher frequencies, it exhibited an exponential decay that was consistent across languages, with individual variations between 80 and 180 Hz, likely reflecting speakers’ fundamental frequencies (f_0_; Figure 1a). Notably, there was no power increase in the low-gamma range (30-50Hz), but a decrease in the exponential decay in the 50-150Hz range (Figure 1a). To estimate whether the speech envelope exhibits cross-frequency interactions, we computed phase-amplitude coupling (PAC; Figures 1b; Figure S1). We identified significant coupling between the primary speech rate (i.e., the phase at ∼2–6 Hz) and amplitudes at ∼30–50 Hz and ∼100–150 Hz (p<.05, surrogate tests based on temporal shifting of the amplitude signal), corresponding to theta, low-gamma, and high-gamma bands (Figure 1c). This pattern was present across individual languages but was absent in musical and environmental sounds (Figure S2), suggesting that the pattern reflects universal features specific to speech. In addition, coupling in the 100–150 Hz range varied across languages and the amplitude frequency exhibiting maximal (and significant) PAC correlated with the f_0_ of each speaker (Pearson correlation across languages; N=11; 6 languages showed no significant high-frequency (80-150 Hz) PAC: R=0.92, df=9, p<0.001; Figure 1d).

**Figure 1:**
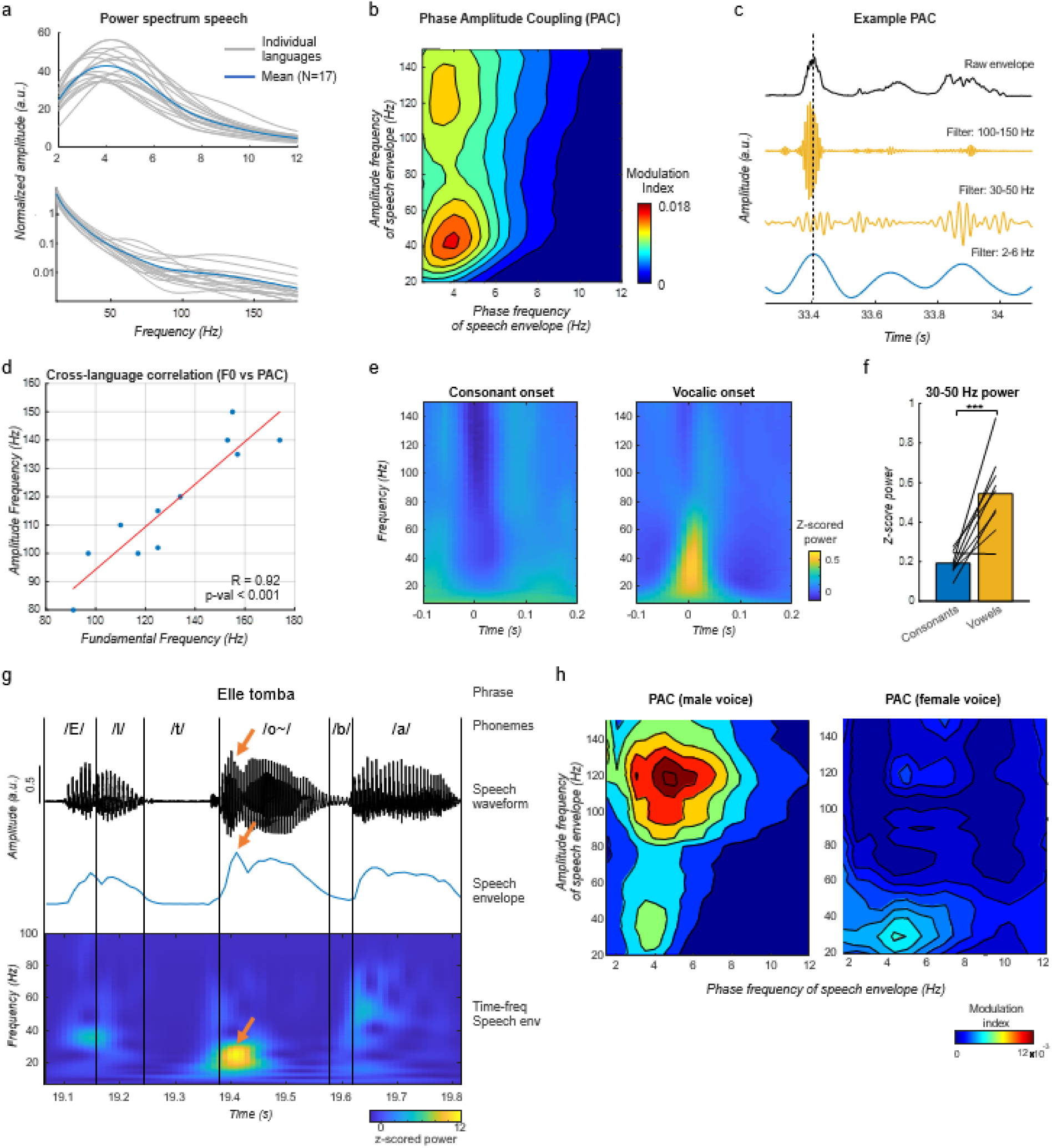
Acoustic structure of the speech envelope across 17 languages. **a)** Power spectrum between 2 and 12 Hz (top) and between 12 and 180 Hz (bottom) of the envelope of naturalistic, discourse-level speech, displayed for each language (gray lines) and averaged across languages (blue line). **b)** Significant phase amplitude coupling (PAC) comodulogram estimated across languages (N = 17), showing coupling between the phase (2-12 Hz) and amplitude (20-150 Hz) of the speech envelope (p<.05, surrogate tests). **c)** Example of speech envelope segment exhibiting strong PAC. From top to bottom: raw envelope, the same signal filtered in the 100–150 Hz, 30–50 Hz, and 2–6 Hz bands. The dashed line indicates the peak of the slow-frequency cycle, aligned with increased amplitude in the higher-frequency bands. **d)** Correlation across languages between the median fundamental frequency (f_0_) and the amplitude frequency exhibiting maximal and significant PAC (between 80 and 150 Hz). **e)** Averaged time-frequency response of the speech envelope aligned to consonant (left) or vocalic (right) onsets, extracted from 9 languages. **f)** Maximum of 30-50 Hz activity (N=9 languages) after phoneme onset for consonants (blue) and vowels (yellow). Black lines represent individual languages (paired t-test, *** p<.001). **g)** Example phrase exhibiting strong 30–50 Hz activity after vocalic onset. From top to bottom: sentence in French, phonetic transcription with onset-aligned phonemes, waveform, envelope, and time-frequency representation of the envelope. Arrows indicate fluctuations in the envelope generating the 30–50 Hz activity. **h)** Significant PAC comodulogram of the speech envelope of the 10-minute French stories used during neural recordings, uttered by (left) a male voice (median f_0_: 139 Hz) and (right) a female voice (median f_0_: 189 Hz; p<.05, surrogate tests).

To obtain further insights into the origin of the 30-50 Hz activity, we analyzed the time-frequency response of the speech envelope locked to phoneme onsets in 9 languages (Figure 1e; Table S1; see Methods). The analysis revealed a 30–50 Hz response related to vocalic onsets —but not seen in in consonants (paired t-test, maximum of 30-50 Hz activity after phoneme onset for consonants vs vowels: t(8)=5.15, p<.001; Figure 1f)— emerging ∼15ms after vocalic onset. Importantly, this activity reflected a transient amplitude increase, lasting ∼25 ms, rather than sustained oscillations (Figure 1g).

We next analyzed the two audio sets used during the SEEG recordings, each consisting of a 10-minute French story narrated by either a male or a female voice (see Methods). In both cases, the speech envelope was characterized by a consistent significant 30-50 Hz amplitude coupling with the primary speech rate (∼5 Hz; p<.05, surrogate tests, Figure 1h). The male voice showed additional coupling in the 100–150 Hz range, consistent with its f_0_ (median f_0_: male voice: 139 Hz; female voice: 189 Hz). In both recordings, gamma-like components were maximal at peak phase (∼0°) of the slow dynamics, indicating alignment with the maximum intensity of the primary speech rhythm (Figure S3).

### The auditory cortex is dominated by alpha oscillations at rest, while theta and gamma dynamics mirror the speech envelope

SEEG activity was recorded from the auditory cortex of 18 epileptic patients (Figures 2a-b) during rest and passive listening to a 10-minute story (male voice; see Methods). Recordings covered medial, primary (TE1.0, TE1.2), and lateral higher-level (Superior Temporal Gyrus (STG); BA41/A42, BA22) auditory areas (Fan et al., 2016). Data from 23 electrodes (5 bilateral implantations; 305 channels) were analyzed using independent component analysis (ICA) to separate neural sources (SEEG-ICs; Figures 2c-d; Herreras et al., 2015; López-Madrona, Trébuchon, Bénar, et al., 2024; López-Madrona, Trébuchon, Mindruta, et al., 2024; Michelmann et al., 2018).

**Figure 2:**
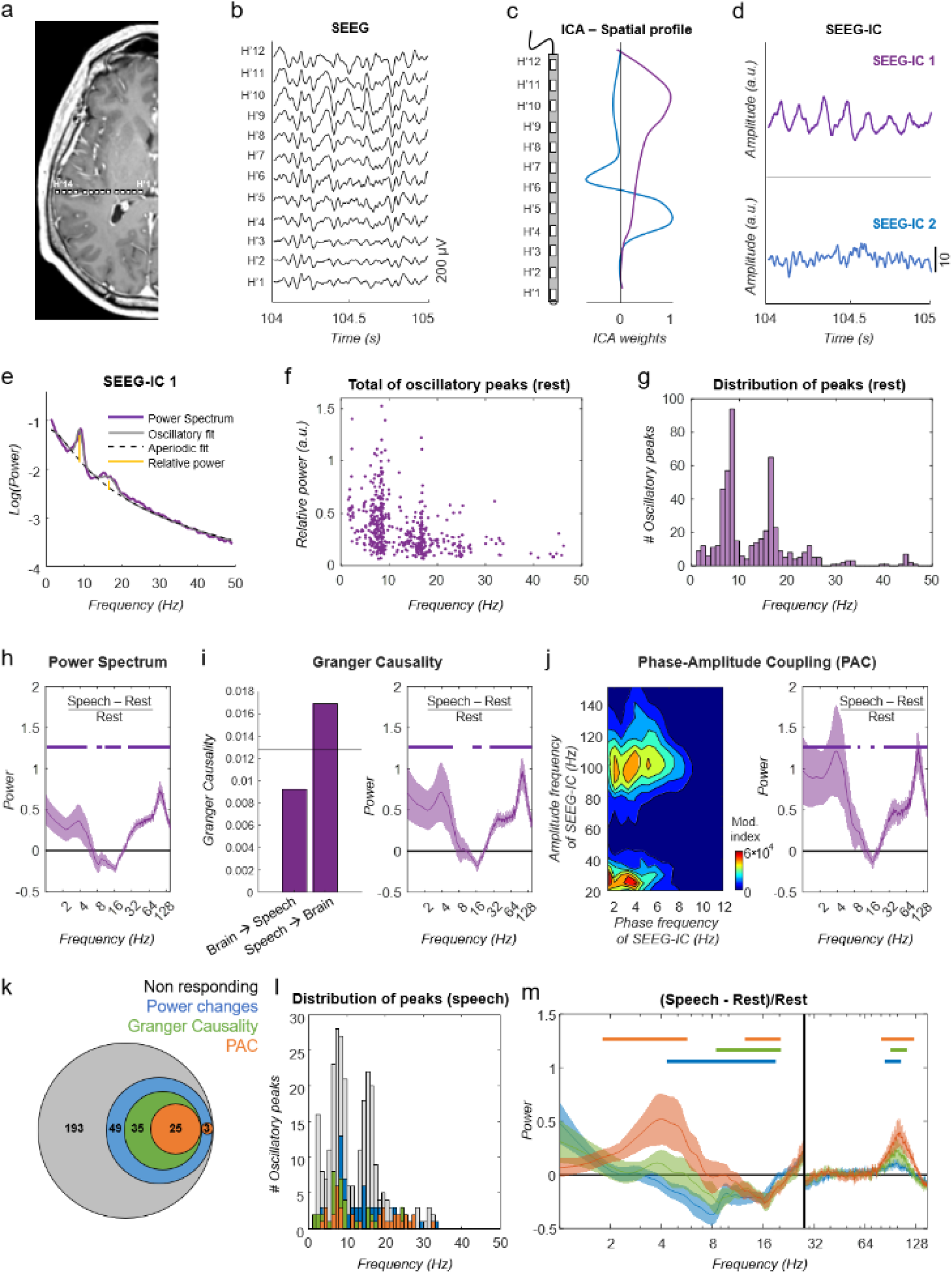
Identification of oscillatory and speech related neural sources. **a-e) Methods. a)** Example of cerebral MRI scan (3D T1-weighted): axial cross section with reconstruction of the SEEG electrode position for patient #6. The location of each contact is represented with white rectangles. **b)** Example of monopolar recordings during resting state. **c)** Spatial profile of two SEEG-ICs (independent components) across the electrode shaft, representing their contribution to each contact. Both components have clear peaks (ICA weights) along the electrode, indicating the origin of the neural activity around these contacts. **d)** SEEG-IC traces during the same time period as panel (b). The first component (SEEG-IC 1) captures high-amplitude low-frequency oscillations from the contacts close to the surface, while SEEG-IC 2 is characterized by irregular activity, with no clear oscillatory pattern. **e)** Schematic of the approach to detect oscillations for SEEG-IC 1. The power spectrum is modeled as a combination of an aperiodic fit and oscillatory peaks. The relative power (yellow lines) of each oscillatory peak is measured as the difference between oscillatory (gray) and aperiodic (dashed) fits. **f-g) Resting state. f)** Relative power of all significant oscillatory peaks identified across patients during resting state. Points represent oscillatory peaks (N=507). **g)** Histogram of frequencies for all the oscillatory peaks in panel (f). **h-m) Speech processing. h)** Normalized power spectra (speech-rest)/rest, averaged across all components exhibiting significant effect (N=112). For all panels: bold lines represent power ratios significantly different from zero at the group level (p<.05, t-tests against zero, FDR-corrected). Error bars are s.e.m. **i)** Left: example of SEEG-IC with a significant Granger Causality (GC) from the speech envelope, but not in the opposite direction (black line represents statistical threshold: p=.05, surrogate tests). Right: Normalized power spectra across all components with a significant GC from the speech envelope (N=60). **j)** Left: example of one SEEG-IC with a significant phase-amplitude coupling (PAC) between the phase of theta and the gamma amplitude (p<.05, surrogate tests). Right: Normalized power spectra across all components with a significant theta-gamma PAC (N=28). **k)** Venn diagram of SEEG-ICs classified by their outcomes in the different analyses: non-responsive channels (N=193; in grey), significant power changes only (N=49; in blue), significant GC and power changes (N=35; in green), significant PAC, GC and power changes (N=25; in orange), and significant PAC and power changes (N=3; in orange). **l)** Histogram of oscillatory peaks during speech processing, for the different groups of SEEG-ICs identified in panel (k). Each group was considered independently, excluding those contained within smaller circles (e.g., the blue histogram represents the SEEG-ICs with significant power changes but no GC or PAC). **m)** Normalized power spectra for the different groups of SEEG-ICs.Aperiodic activity was corrected, with different fits between 1–28 Hz and 28–150 Hz (vertical black line).

Spectral analysis of SEEG-ICs at rest indicated a combination of aperiodic background activity and oscillatory components (Donoghue et al., 2020; Figure 2e). Most SEEG-ICs exhibited prominent alpha-like oscillations (6–9 Hz, N=191, Figures 2f-g; López-Madrona et al., 2024), along with their harmonics (12–18 Hz, N=151). Delta activity appeared in a subset of the data (∼2 Hz, N=21), and remaining peaks outside these ranges were sparse and of low power. Resting state activity was notably characterized by an absence of oscillatory activity at theta and/or low-gamma rates.

To characterize neural responses to speech, we applied three complementary analyses. First, we examined spectral power changes during speech processing compared to rest. Significant power increases were observed in the delta/theta (1–6 Hz) and broadband gamma (30–150 Hz) ranges across ∼37% of SEEG-ICs (N=112), alongside suppression in the alpha/beta band (6–20 Hz; p<.05, (speech-rest)/rest ratio: individual permutation tests, FDR-corrected; Figure 2h; non-normalized spectra in Figure S4).

Second, we computed speech-brain (temporal) Granger Causality (GC) and found significant GC from the speech envelope in 60 SEEG-ICs, meaning speech significantly predicted brain activity (speech-driven SEEG-IC), with none showing the opposite pattern (p<.05, surrogate tests based on temporally shifted speech; Figures 2i and S4). All these speech-driven sources were also characterized by a significant power increase (p<.05, individual permutation tests), particularly in the delta/theta (1–6 Hz) and broadband gamma (30–150 Hz) ranges. Additionally, 37/60 SEEG-ICs showed reduced alpha/beta power (6–20 Hz; p<.05, individual permutation tests).

Finally, we analysed neural phase-amplitude coupling (PAC) and observed significant PAC between theta (peaking at ∼4 Hz) and gamma dynamics in 28 SEEG-ICs (p<.05, surrogate tests based on temporal shifting of the amplitude signal; Figures 2j and S4). Two distinct gamma bands were coupled with theta: a high-gamma component (peaking at ∼110 Hz) in all 28 SEEG-ICs, and a low-gamma component (peaking at ∼40 Hz) in 12 of them. The 28 SEEG-ICs also showed increased delta/theta (1–6 Hz) and broadband gamma (30–150 Hz) power (p<.05, individual permutation tests), with 25/28 also showing significant GC from the speech envelope, indicating direct modulation by speech.

Importantly, the three analytic approaches ranged from least to most selective: power change, GC, and PAC (Figure 2k). We thus classified SEEG-ICs into three groups: power changes only (N=49), GC without PAC (N=35), and PAC (N=28). Only 3 channels did not entirely fit this nested structure (i.e., showing significant PAC (and power change), but not GC, see above). As in the resting condition, all groups exhibited a dominant alpha-like rhythm and its first harmonic (Figure 2l). However, speech elicited additional, group-specific changes in power at other frequencies (Figure 2m). The *PAC group* was the only one with a significant theta power increase (p<.05 group level t-tests between speech and rest, FDR-corrected; Figure 2m). All groups exhibited increased high-gamma power (∼110 Hz) and beta suppression (12–20 Hz). High-gamma enhancement peaked in the *PAC group*, while beta suppression was strongest in the *power changes group* and extended into the alpha range (∼8 Hz), the dominant resting-state rhythm (see Figure 2g). Overall, these findings indicate that theta and gamma neural oscillations are directly driven by the speech envelope, inducing PAC between these rhythms in the corresponding neural sources.

### Theta and gamma responses are spatially segregated in the auditory cortex

To further characterize how speech drives theta and gamma responses in the auditory cortex, we computed spectral GC between the speech envelope and the speech-driven SEEG-ICs (i.e., with significant temporal GC, N=60; see Figure 2i). Three main speech-driven frequencies emerged: theta (∼2–6 Hz), low-gamma (∼25–50 Hz), and high-gamma (∼100–150 Hz), with SEEG-ICs exhibiting one or more of these dynamics (Figure 3a). Three profiles (groups) of SEEG-ICs were differentiated, with 10 SEEG-ICs modulated only at theta, 26 only at (low and/or high) gamma, and 24 at both frequencies.

**Figure 3:**
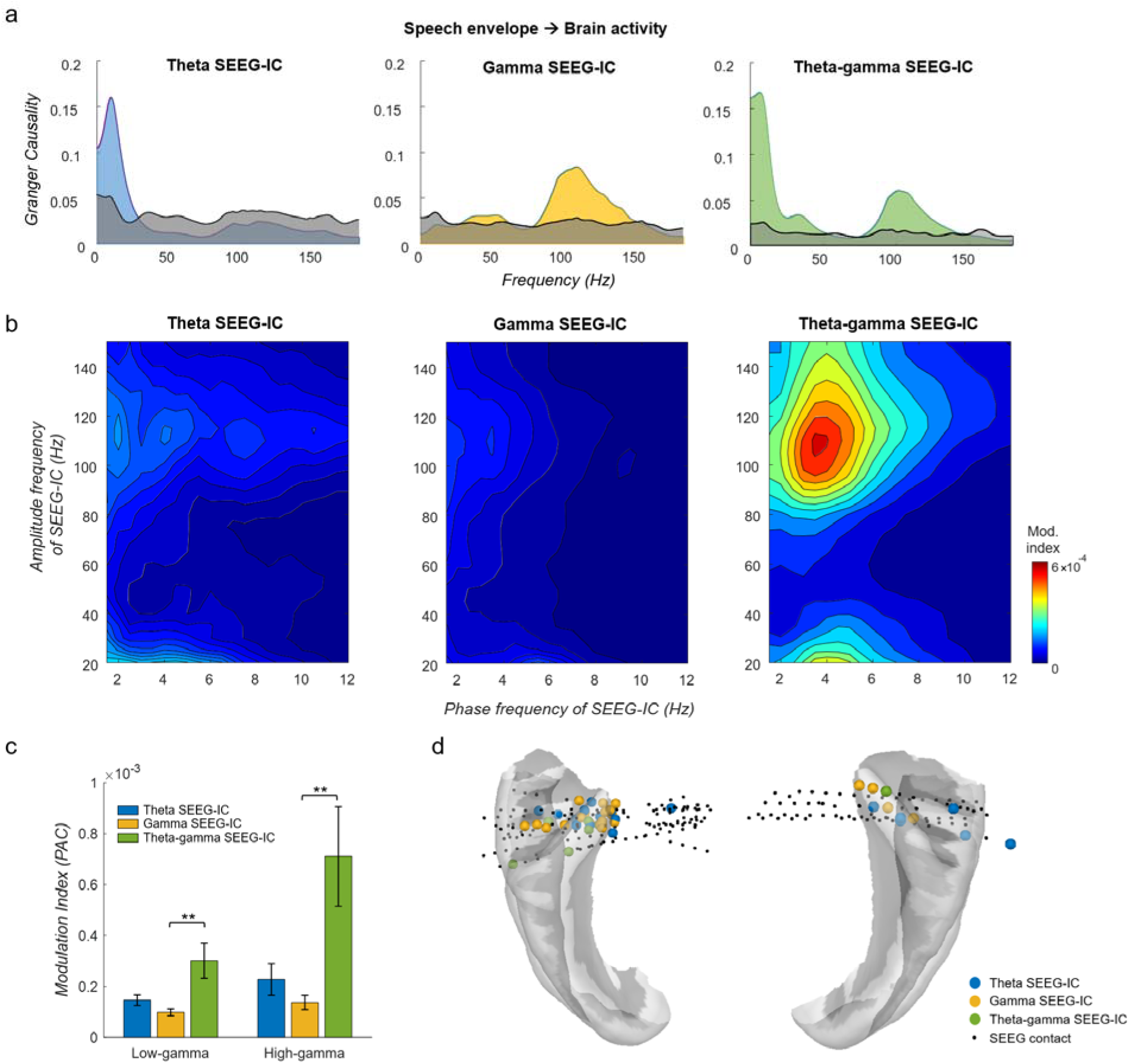
Spectral GC and neural PAC profiles. **a)** Representative example of the three types of outcomes observed in the spectral GC analysis between the speech envelope and neural activity: theta (2–6 Hz; in blue), gamma (low 30–50 Hz and/or high 100–150 Hz; in yellow), or theta and gamma (low and/or high; in green) significant GC. Shaded areas indicate the significance threshold (surrogate tests). **b)** Average PAC modulation spectrum for the three groups of SEEG-ICs: theta (left; N=10), gamma (middle; N=26) and theta-gamma (right; N=24). **c)** PAC between the theta (2–6 Hz) phase and low- (30–50 Hz) or high-gamma (100–150 Hz) for each SEEG-IC group (** p<.05, Tukey’s HSD post-hoc test). **d)** Position of the 60 SEEG-ICs from the three groups on a 3D surface of the temporal lobe. The location of each SEEG-IC is defined by the contact with the maximal contribution in the spatial IC profile.

This last group of SEEG-ICs was characterized by stronger PAC (Figure 3b), at both low-gamma (one-way ANOVA: F(2,57)=5.4, p=.007; Tukey’s HSD: gamma vs. theta-gamma, p=.006; all other comparisons: p>.16) and high-gamma frequencies (one-way ANOVA: F(2,57)=5.7, p=.005; Tukey’s HSD: gamma vs. theta-gamma, p=.005; all other comparisons p>.10; Figure 3c). This indicates that theta and gamma dynamics —and their interaction (PAC)— are directly driven by the speech envelope.

Finally, as SEEG-ICs correspond to distinct neural sources, these three groups were spatially segregated along the auditory cortex (Figure 3d), mainly laterally, within area 41/42 of the superior temporal gyrus (STG), though no specific region across participants was associated with a particular dynamics (Table S2). This shows that theta and gamma responses occur in overlapping anatomical regions but are generated by distinct neural populations (SEEG-IC), indicative of a functional segregation of these speech-driven neural dynamics.

### High-gamma (100-150 Hz) response is a mixture of frequency-following and evoked activity

While theta and low-gamma activity during speech processing are driven by the speech signal, high-gamma (100-150 Hz) activity exhibits a more complex response profile. It peaked in SEEG-ICs with significant PAC, but was also present in SEEG-ICs showing only power changes —i.e., without significant speech-brain GC (Figure 2m). This indicates that a large portion —but not all— of the activity in this frequency range reflects the frequency-following response to the fundamental frequency of the male voice used in our primary stimulus (f_0_ = 139 Hz; Figure 1h, left panel). To dissociate these two sources of high-gamma activity, we recorded a second cohort of patients (N=2) during passive listening to the same male-narrated story and another story narrated by a female voice (f₀ = 189 Hz; Figure 1h right panel; see Methods).

We identified three speech-driven SEEG-ICs (N=3) with significant temporal GC in both auditory conditions. Two were classified as theta SEEG-IC, showing consistent modulation at the speech rate across stimuli (Figure 4a, top). The third was a gamma SEEG-IC (Figure 4a, bottom), exhibiting low-gamma modulation in both recordings. High-gamma modulation was also observed, but only for the male voice, consistent with a frequency-following response to f_0_ (see Figure 1h).

**Figure 4:**
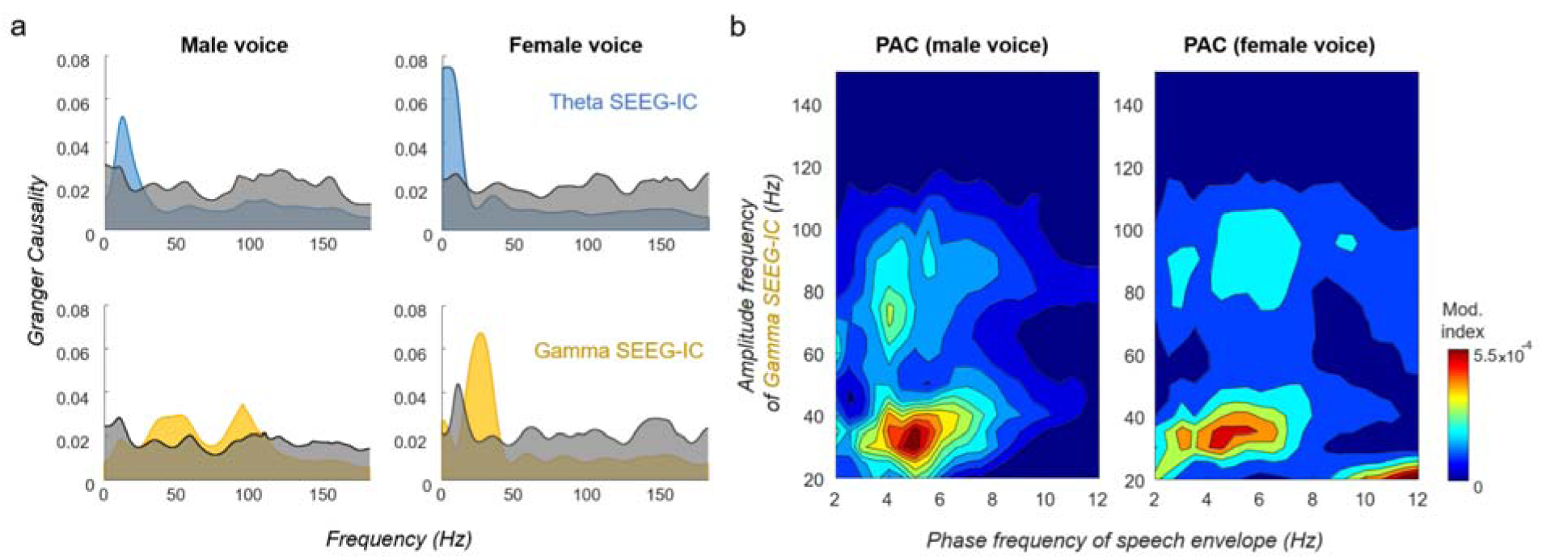
Relation between speech-brain coupling and fundamental frequency. **a)** Comparison of neural responses to male and female recordings with representative examples of the two types of outcomes observed in the spectral GC analysis between the speech envelope (left: male voice; right: female voice) and neural activity: theta (2–6 Hz; top; in blue) and gamma (low 30–50 Hz and/or high 100–150 Hz; bottom; in yellow). Shaded areas indicate the significance threshold (surrogate tests). **b)** PAC modulation spectrum estimated between the phase of the speech envelope (male: left, female: right) and the amplitude of the gamma SEEG-IC from panel (e) (p<.05, surrogate tests).

We next analysed speech-brain PAC in the gamma SEEG-IC (Figure 4b). In both recordings, gamma amplitude was modulated at the speech rate (∼5 Hz), with dominant coupling in the low-gamma band (30–50 Hz), both for male and female voices. High-gamma PAC (100–150 Hz) was also significant in both conditions (p<.05, surrogate tests based on temporal shifting of the SEEG-IC), indicating speech-related high-gamma activity independent of f_0_. These results show that while high-gamma reflects an intrinsic neural rhythm, it can also be linearly driven by speech acoustics when the fundamental frequency falls within its frequency range.

### Gamma to theta PAC in the auditory cortex

While theta and gamma neural dynamics primarily mirror speech acoustics, they are spatially segregated in the auditory cortex (Figure 3d). To investigate directed functional connectivity between the speech-driven SEEG-ICs (i.e., with significant temporal GC, N=60), we computed the cross-frequency directionality (CFD) index. We first computed the CFD on the speech envelope and found no significant directionality (all p>.05, surrogate tests based on temporal shifting of the amplitude signal), indicating that slow (∼2–6 Hz) and fast (∼30–150 Hz) envelope dynamics evolve overall synchronously.

Then, we estimated the speech-brain CFD comodulogram across all speech-driven SEEG-ICs with significant gamma modulation (N=50; see Figure 3a), and confirmed that the main speech rate (∼4 Hz) primarily modulates low-gamma (30-50 Hz; one-sample t-test against zero: t(49)=6.5, p<.001) and high-gamma (100-150 Hz; t(49)=6.8, p<.001) neural activity (Figure 5a). Additionally, analysis of the speech-driven SEEG-ICs with significant theta modulation (N=34; see Figure 3a) showed that neural theta activity (∼4 Hz) is primarily modulated by acoustic dynamics of the speech envelope at 30-50 Hz (t(33)=-8.82, p<.001) and 100-150 Hz (t(33)=-10.57, p<.001, Figure 5b). As slow (∼4 Hz) and fast (∼30–50 and ∼100-150 Hz) dynamics are strongly coupled in the speech envelope, this result corroborates the spectral GC analysis, which already revealed linear, directed speech-brain functional connectivity at theta, low-gamma, and high-gamma frequencies (Figure 3a).

**Figure 5:**
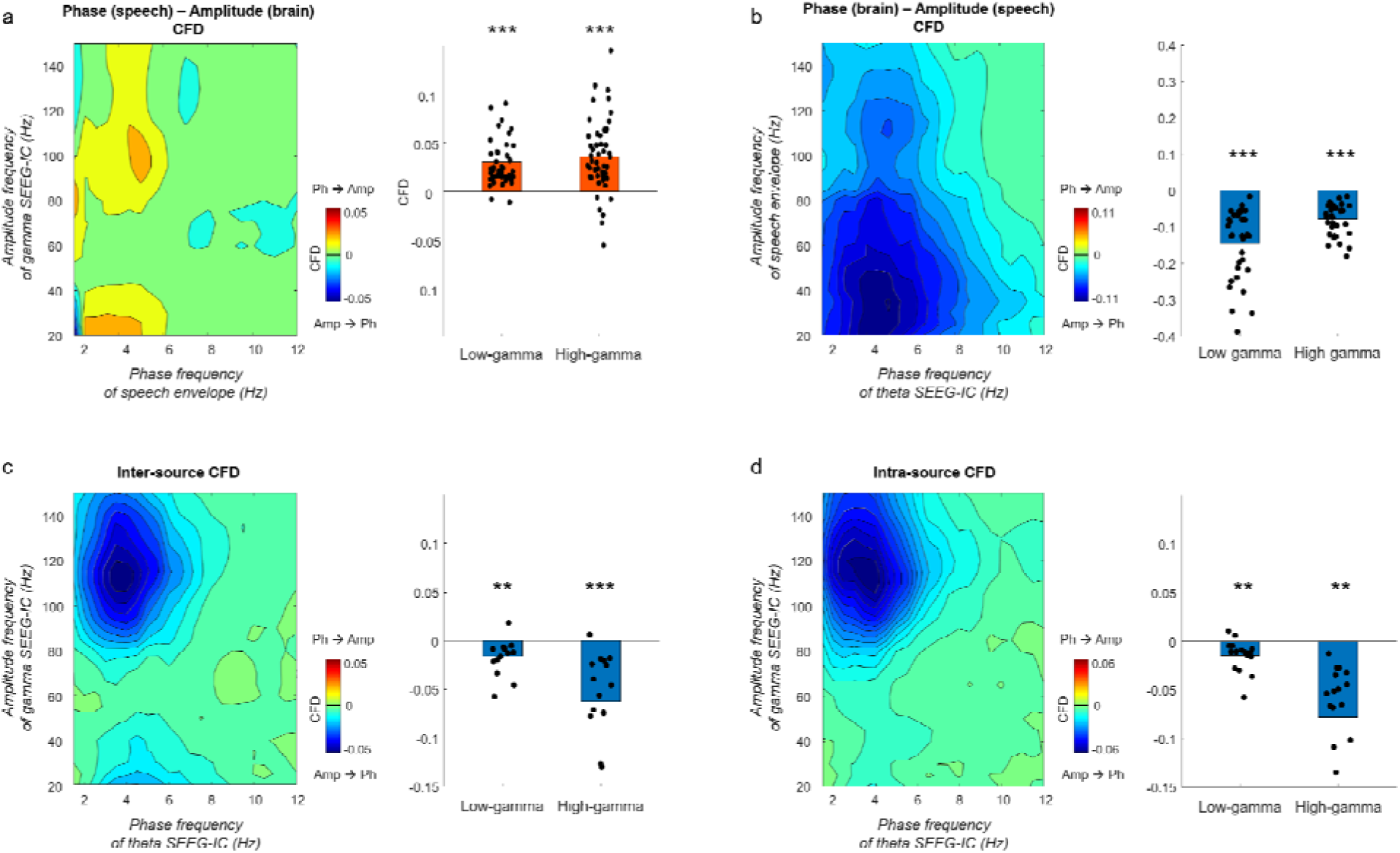
Directed speech-brain and brain-brain PAC. **a)** Left: Averaged speech-brain cross-frequency directionality (CFD) comodulogram, estimated between the phase of the speech envelope and the amplitude of speech-driven gamma SEEG-ICs (N=50). Right: Average CFD between theta phase (2-6 Hz) and low- (30-50 Hz) or high-gamma (100-150 Hz) amplitude. **b)** speech-brain CFD estimated between the phase of speech-driven theta SEEG-ICs (N=34) and the amplitude of the speech envelope. **c)** Inter-source CFD averaged across electrodes (N=15). For each electrode a single CFD map was computed as the average of all its speech-driven pairs. **d)** Intra-source CFD averaged across electrodes (N=16). Points represent (a,b) individual SEEG-ICs or (c,d) electrodes. Positive CFD values indicate phase-to-amplitude directionality; negative values indicate the reverse. One-sample t-test against zero:: ** p<.01, *** p<.001.

We then assessed functional directionality between neural sources by computing CFD between all pairs of speech-driven SEEG-ICs (N=60; see Figure 2i) per electrode (N=15; 8 electrodes were excluded due to having N < 2 significant speech-driven SEEG-ICs). CFD maps were averaged within each electrode shaft, excluding within-source analysis. Surprisingly, gamma amplitude consistently drove theta phase across electrodes, for both low-gamma (t(14)=-3.36, p=.005) and high-gamma (t(14)=-4.17, p<.001) activity (Figure 5c). Finally, intra-source CFD within theta–gamma SEEG-ICs (averaged CFD for each electrode with theta-gamma SEEG-ICs, N=16) revealed the same pattern: gamma amplitude consistently drove theta phase, for both low-gamma (t(15)=-3.57, p=.003) and high-gamma (t(15)=-3.72, p=.002) activity (Figure 5d). Together, these results show that gamma activity—both low and high— drives theta phase, both within and across neural sources, suggesting that auditory theta activity is shaped by gamma activity, rather than the reverse.

We finally estimated the latency of the neural responses, by computing frequency-resolved linear speech-brain cross-correlation analyses (Figure S5). For each speech-driven SEEG-IC (N=60), we computed the lag between the speech envelope and (i) the theta-filtered neural activity (2–6 Hz) for theta SEEG-ICs (N=33 SEEG-ICs showing a significant cross-correlation with the speech envelope, see Methods), and (ii) the envelope of low- (30–50 Hz; N=22) or high-gamma (100–150 Hz; N=29) signals for gamma GC SEEG-ICs. Delays in theta SEEG-ICs were highly variable (range: 8–205 ms; mean ± s.d.: 78.2 ± 63.6 ms). In contrast, delays were more consistent in the gamma bands (low-gamma: 58.8 ± 9.4 ms; high-gamma: 23.2 ± 33.2 ms). High-gamma responses occurred significantly earlier than both theta (one-way ANOVA: F(2,81)=11.82, p<.001; Tukey’s HSD: theta vs. high-gamma, p<0.001) and low-gamma (p=.017), with no significant difference between theta and low-gamma responses (p=.26), indicating that higher frequency activity is processed earliest in the auditory cortex.

## DISCUSSION

### Acoustic origin of theta and gamma dynamics in early auditory cortex

This study demonstrates that multiplexed theta–gamma dynamics in the human auditory cortex during speech perception are not reflective of an intrinsic neural code of the auditory cortex but mirror acoustic features present in the speech signal (Figure 6). As a highly evolved human signal, these features are likely related to the processing capacities of the auditory cortex, without relying on neural oscillations present at rest. Using SEEG recordings, we identified multiple auditory neural sources that are selectively activated by natural speech and remain silent at rest (Figure 2). Each source is driven by frequency-specific components of the speech envelope that approximate distinct linguistic features and are robust across languages: syllabic rate (theta), vowel energy (low-gamma), and/or fundamental frequency (f_0_; high-gamma; Figure 1 and S1). These acoustic features are specific to speech and absent in musical and environmental sounds (Figure S2), underscoring the unique temporal structure of speech. Crucially, phase-amplitude coupling (PAC) between these frequencies —often interpreted as a mechanism for hierarchical parsing— is also present in the speech signal itself, reflecting its nested acoustic architecture (Figure 1). Each speech-driven source within the same auditory region is feature-selective, indicating functional segregation (i.e., demultiplexing; Figure 3). They are functionally connected via directed PAC, with gamma activity —both frequency-following and evoked (Figure 4)—driving theta phase (Figure 5), a reversal of the dominant model. Overall, these results challenge the interpretation of auditory cortical PAC as an endogenous mechanism for speech parsing.

**Figure 6:**
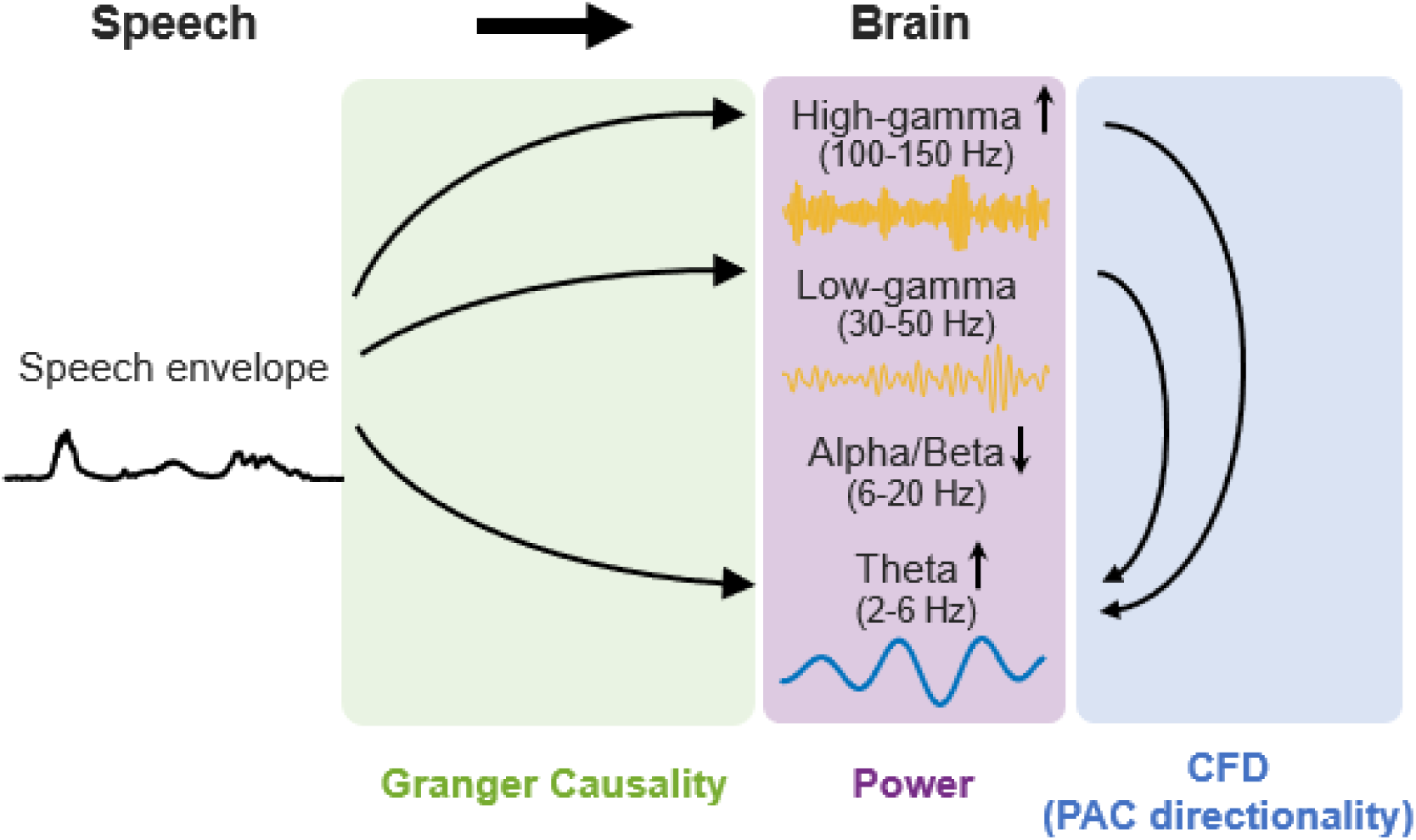
Diagram summarizing the relationship between the speech signal and early auditory cortex dynamics during speech perception. The green box (Granger Causality) shows brain dynamics that mirror the speech envelope, linearly driven by it, indicated by directional arrows. In the purple box (Power), up/down vertical arrows reflect increases or decreases in power during speech perception compared to rest across all neural sources (SEEG-ICs). The blue box (Cross-Frequency Directionality, CFD) depicts internal connectivity between brain dynamics associated with phase–amplitude coupling (PAC).

### From temporal hierarchies to parallel multiplexing

Speech encodes multiple layers of information concurrently (Coupé et al., 2019; Rosen, 1992)—such as speaker identity, pitch (f₀), timbre, prosody, semantics, and syntax. These layers have been shaped by evolution, from the cycles of mandibular oscillation in the articulatory system to the development of motor control circuits in the brain (Fitch, 2000; MacNeilage, 1998). This has resulted in dynamical patterns unique to human speech. While linguistic theory has long described a hierarchical organization of units (features, phonemes, syllables, morphemes, prosodic words, phrases, sentences), whether these are systematically embedded in a dynamically structured acoustic signal has not been quantitatively assessed.

Brain rhythms are also hierarchically organized, aligning excitability across timescales to optimize coding efficiency (Bragin et al., 1995; Buzsáki & Draguhn, 2004; Freeman & Rogers, 2002; Lakatos et al., 2005; Lisman & Idiart, 1995; Pöppel, 1997; Steriade, 2006). In particular, the theta-gamma neural code (Canolty & Knight, 2010; Lisman & Jensen, 2013) which —though primarily studied in the hippocampus— proposes a general encoding mechanism, whereby theta oscillations structure the timing of gamma bursts, representing discrete units of information.

Building on the isomorphism between brain rhythms and speech features (e.g., theta/syllables, gamma/phonetic features; Poeppel, 2001, 2003), Giraud & Poeppel, (2012) extended this framework to speech, proposing that such correspondence supports the chunking and tracking of speech units in time. The nested alignment of fast (gamma) and slow (theta) cortical rhythms has been widely observed during speech processing (Attaheri et al., 2022; Gross et al., 2013; Hyafil, Fontolan, et al., 2015; Koutsoukos et al., 2013; Lizarazu et al., 2019, 2023; Mai et al., 2016; Meyer et al., 2020; Morillon et al., 2012; J. Wang et al., 2014). Such multiplexed temporal coding has gained support as a biologically plausible mechanism for mapping speech’s multiscale structure onto neural dynamics. By shifting focus from categorical linguistic units toward the temporal dimension (Norman-Haignere et al., 2025), it offers a unified framework in which hierarchically embedded speech units are organized within a shared temporal architecture.

However, the common temporal structure of speech and brain rhythms raises a critical question: does phase-amplitude coupling in the auditory cortex emerge de novo in the early cortical stages of speech processing for re-mapping speech from acoustics to other representational forms suited to neural processing, as proposed in other cognitive functions, or is it evoked by the temporal structure of the acoustic signal itself?

### Revisiting the role of PAC in speech processing

This question has not been systematically, quantitatively, and mechanistically examined, and the field has largely assumed a neural origin. However, our findings reveal that PAC also characterizes the speech envelope itself, exposing cross-frequency structure not visible in the power spectrum alone (Figure 1a-b). This suggests that the nested dynamics often attributed to endogenous auditory parsing in fact arise from the stimulus.

While our results are compatible with the multiplexing described by the multi-timescale hypothesis (Baroni et al., 2020; Giraud & Poeppel, 2012; Giroud et al., 2020; Teng et al., 2016, 2017; Teng & Poeppel, 2020), they challenge the assumption that nested brain rhythms reflect endogenously generated parsing mechanisms in early auditory cortex. Here, we show that theta–gamma coupling is explainable by the speech acoustic features, and that the early auditory cortex *mirror*s a highly structured input rather than generates internal windows for segmentation. This reframes the idea that the auditory cortex operates on multiple timescales: theta, low-gamma, and high-gamma reflect syllabic rate, vowel onsets, and f₀, respectively (Giroud et al., 2024; Kulasingham et al., 2020). These features are already dissociable in the stimulus and processed by distinct but spatially overlapping neural populations, supporting the idea of parallel streams for spectrotemporal parsing (Poeppel, 2003). This challenges the prevailing view that PAC in the auditory cortex supports endogenous hierarchical parsing —as observed in the hippocampus (Bragin et al., 1995; Colgin et al., 2009; López-Madrona et al., 2020)— and instead suggests that it reflects the brain’s alignment to a temporal code already embedded in speech.

An important finding is that we did not detect oscillatory activity at rest, which is supported by the lack of consensus in the literature regarding intrinsic auditory rhythms. Across studies, proposed resting-state cortical auditory oscillations span a wide frequency range: delta (Armonaite et al., 2022; Frauscher et al., 2018; Groppe et al., 2013; Kalamangalam et al., 2020; Mellem et al., 2017), theta (Giraud et al., 2007; Morillon et al., 2012), beta (Ramkumar et al., 2014), or low-gamma (Giraud et al., 2007; Groppe et al., 2013; Hillebrand et al., 2012; Morillon et al., 2010). Some studies report endogenous activity across nearly all frequency bands (Keitel & Gross, 2016; Lubinus et al., 2021; Neymotin et al., 2022). However, the most consistently reported and replicable finding across studies is a prominent alpha-like peak —often referred to as the auditory “tau” rhythm (Capilla et al., 2022; Frauscher et al., 2018; Groppe et al., 2013; Hillebrand et al., 2012; Kalamangalam et al., 2020; Keitel & Gross, 2016; Lehtelä et al., 1997; López-Madrona, Trébuchon, Bénar, et al., 2024; Mahjoory et al., 2020; Niedermeyer, 1990; Ramkumar et al., 2014; Tiihonen et al., 1991) — in line with our own observations (Figure 2). This variability, along with the methodological difficulty of disentangling true oscillatory peaks from broadband 1/f aperiodic activity (Donoghue et al., 2020), casts doubt on the notion that theta and gamma reflect stable intrinsic rhythms in the human auditory system.

However, the absence of prominent oscillatory activity at rest does not imply a lack of resonance properties. Instead, latent capacities of the auditory cortex may manifest as a preferential sensitivity to specific frequencies during auditory stimulation. This is the case of auditory steady-state responses (ASSRs), in which cortical circuits preferentially entrain to periodic auditory inputs typically around 40 Hz (Arnal et al., 2019; Galambos et al., 1981; Mäkelä & Hari, 1987; Picton et al., 2003). ASSRs are not just a series of discrete evoked potentials, but rather a response to the periodicity of the stimulus, where the phase of neural activity locks to the stimulus modulation frequency. They illustrate the resonance properties of auditory circuits —namely, the frequency preferences of neural populations shaped by neural connectivity and synaptic time constants (Hutcheon & Yarom, 2000). Thus, resonance differs from spontaneous oscillations in that it does not reflect the system’s intrinsic dynamics, but rather its capacity to preferentially respond to external input at specific frequencies (Giroud et al., 2020; Teng et al., 2016, 2017; Teng & Poeppel, 2020, Lehongre et al.,2011).

In this context, the observed mirroring of both low- and high-gamma speech features by auditory cortex aligns with findings from the frequency-following response (FFR) literature. FFRs —traditionally measured in subcortical structures but increasingly documented cortically— reflect phase-locked responses to periodic stimulus features such as the fundamental frequency or high-rate temporal modulations (Bidelman, 2018; Coffey et al., 2019; Nourski, 2017). Importantly, intracranial and high-resolution imaging data demonstrate that human auditory cortex can phase-lock to acoustic frequencies up to ∼130 Hz (Brugge et al., 2009; Nourski & Brugge, 2011; Steinschneider et al., 2013), overlapping with the gamma-band responses we identify here. This supports the interpretation that both low- and high-gamma PAC in the auditory cortex reflect frequency-following responses to acoustic modulations, rather than internally generated sampling mechanisms (Lerousseau et al., 2021). Moreover, the distinct spatial patterns of gamma and theta SEEG sources are consistent with the spatial organization of FFRs and temporal envelope tracking described in auditory cortex (Nourski, 2017), suggesting that different neuronal ensembles are tuned to resonate to specific spectrotemporal features of speech.

More broadly, phase–amplitude coupling is a well-established organizational principle in the brain, supporting the compression and multiplexing of information across timescales (Akam & Kullmann, 2012; Canolty et al., 2006; Hyafil, Fontolan, et al., 2015). Previous hypotheses have adapted this mechanism to the context of speech processing. Accordingly, PAC would align perception with the richly structured input while operating within the temporal resolution and bandwidth constraints of cortical circuits. By aligning multiscale acoustic features into dedicated oscillatory neural channels, PAC may enhance both perceptual resolution and neural efficiency — a principle that underpins computational models designed to match neural auditory dynamics (Baroni et al., 2020; Dogonasheva et al., 2025; Giraud & Poeppel, 2012; Hovsepyan et al., 2020, 2023; Hyafil, Fontolan, et al., 2015). Our findings question this view as we show that PAC is present in the structure of the speech stimulus itself. Auditory neural PAC may thus not be a mechanism to transform the acoustic structure of speech into linguistic units, but may represent nested temporal regularities already embedded in the signal. This aligns with theoretical work suggesting that the rhythmic structure of speech arises from the biomechanical and cognitive constraints of speech production (O’Dell & Nieminen, 1999; Tilsen, 2016). Rather than generating hierarchical parsing schemes de novo, the auditory cortex may inherit a structured input shaped by the audio-phonological dynamics, mapping it onto internal representations that are both perceivable and reproducible.

We conjecture that PAC reflects the alignment of distributed neural populations to shared temporal anchors. The auditory cortex may thus function as a temporal ‘demultiplexer’, selectively amplifying and routing speech features across oscillatory channels tuned to biologically relevant timescales. This perspective invites a re-evaluation of neural coupling in speech processing. Rather than implementing linguistic parsing, theta–gamma PAC may serve as a biologically efficient mechanism for synchronizing perception to an highly-evolved structured signal. Future work should identify how and where this temporal scaffolding supports the formation of abstract linguistic representations —likely beyond the auditory cortex.

### Directionality of PAC: from gamma to theta

Traditional models of PAC assume that slower rhythms, such as theta, modulate the amplitude of faster gamma activity, creating periodic “windows” of heightened excitability for sensory encoding (Giraud & Poeppel, 2012; Gross et al., 2013; Hyafil, Fontolan, et al., 2015; Lakatos et al., 2008; Lisman & Jensen, 2013; Lizarazu et al., 2019; Mai et al., 2016). In this framework, theta phase gates gamma activity, supporting hierarchical parsing. However, this directionality has rarely been empirically tested, and emerging work has challenged its universality (Baroni et al., 2020; Chalkiadakis et al., 2025; Jiang et al., 2015, 2020; López-Madrona et al., 2020).

Our results provide evidence that low- and high-gamma activity systematically precedes and modulates theta phase, both across distinct SEEG sources and within single neural populations (Figure 5). This gamma-to-theta directionality suggests that fast spectrotemporal features in speech—particularly in the low- and high-gamma ranges—drive early neural responses that in turn align the phase of theta oscillations to incoming input. In this view, gamma responses reflect rapid sensory encoding, while theta dynamics serve to temporally coordinate slower integrative processes, such as syllabic integration.

This finding is consistent with recent proposals of a reversed hierarchy in auditory processing (Baroni et al., 2020; Kösem & van Wassenhove, 2017; Xu et al., 2023), where high-gamma power anticipates theta rhythms and better tracks fast-varying speech features. Our data support this temporal asymmetry, with high-gamma exhibiting shorter latencies than theta (Figure S5), and decoding analyses pointing to a primary auditory origin for gamma responses (Xu et al., 2023). While an alternative explanation could be that theta and gamma are independently entrained by the same stimulus with differing delays, the presence of gamma-to-theta directionality within the same SEEG-ICs (i.e., the same neural source) argues against this account (Hyafil, Giraud, et al., 2015). Taken together, these results call for a re-evaluation of the mechanisms underlying theta–gamma PAC in speech processing, shifting focus from theta-based parsing or segmentation to gamma-driven alignment.

### Roles of gamma-nested dynamics

The 30–50 Hz component of the speech envelope is closely linked to phonemic features and is essential for speech intelligibility (Chait et al., 2015; Shannon et al., 1995). Our findings show that the auditory cortex linearly follows this frequency range (Figure 2a). However, gamma dynamics have also been associated with higher-level linguistic processing and the transformation of acoustic input into linguistic meaning (Di Liberto et al., 2015; Mesgarani et al., 2014). Additionally, while 30–50 Hz energy in the speech envelope is phase-locked to the syllabic rate—a pattern conserved across languages (Figure 4)—the corresponding PAC in the brain is stronger for intelligible than reversed speech (Gross et al., 2013). These findings raise the question of how much gamma activity reflects acoustic encoding versus cognitive processing in the early auditory cortex.

Our data confirm that at least two gamma regimes are at play in auditory cortex during speech processing. The first, captured by our recordings, reflects an early stage of acoustic encoding (Xu et al., 2023). It mirrors low-level features such as phoneme energy and f₀ in a linear, stimulus-driven fashion, and it couples with theta phase at the syllabic rate (Figures 3b and 4). This gamma activity mirrors input dynamics: increasing or decreasing speech rate shifts the gamma frequency proportionally (Lizarazu et al., 2019). In turn, this gamma input may reset the phase of theta oscillations, aligning slower rhythms with syllabic structure. Although this low-gamma regime has an acoustic origin (Howard & Poeppel, 2010), some evidence suggests it may also contribute to speech comprehension. The auditory cortex exhibits a preferential response for 30–50 Hz signals (Teng & Poeppel, 2020), and gamma and theta components often arise from distinct neural populations (Figure 2a). Moreover, gamma activity may be modulated by top-down influences, including delta–gamma coupling (Fontolan et al., 2014). During degraded or unintelligible speech, gamma amplitude and its modulation of theta phase are both reduced (Gross et al., 2013; Lizarazu et al., 2023), impairing theta tracking of the input (Etard & Reichenbach, 2019; Vanthornhout et al., 2018). In kids with autism, theta-gamma coupling is absent and replaced by an atypical beta-gamma coupling, showing that neural coupling phenomena are under neuromodulatory influence (X. Wang et al., 2023).

Beyond this early encoding stage, a second gamma regime may operate in higher-order cortical areas (Di Liberto et al., 2015; Forseth et al., 2020; Hamilton et al., 2018). This activity, which may not couple to theta, could reflect internally generated activity that support higher-level linguistic or semantic operations. Thus, gamma activity in speech processing likely spans a functional continuum —from early sensory tracking to abstract computation— with the presence or absence of PAC distinguishing these stages.

## Conclusion

Our results challenge the prevailing view that theta–gamma phase–amplitude coupling (PAC) in the auditory cortex reflects endogenously generated speech processing mechanisms. Instead, we show that these dynamics largely mirror the temporal structure of the speech signal itself. Theta, low-gamma, and high-gamma activity mirror distinct acoustic features — syllabic rate, vowel-related energy, and f₀— embedded in the speech envelope. This coupling is exogenous, robust across languages, and could be shaped by audio-motor constraints on speech production. This points to a likely universal, biologically rooted mechanism that underpins the human-specific communication niche that is speech.

Crucially, gamma activity leads theta phase, suggesting an active role in aligning slower rhythms with fast-changing acoustic inputs. We demonstrate that these rhythms are processed by distinct but interconnected cortical populations, implying demultiplexing and functional coordination rather than passive responses. While PAC may support speech comprehension, our findings emphasize the need to methodically disentangle intrinsic neural activity from stimulus-driven dynamics. Altogether, these results call for a revised framework: PAC in the auditory cortex mostly reflects the brain’s response to an already-structured signal, not its internal construction.

## ACKNOWLEDGEMENTS

Research supported by grants ANR-16-CONV-0002 (ILCB) and the Excellence Initiative of Aix-University (A*MIDEX AMX-19-IET-004). B.M. is supported by ANR-20-CE28-0007, Fondation Pour l’Audition (FPA RD-2022-09), and co-funded by the European Union (ERC, SPEEDY, ERC-CoG-101043344). L.L. is supported by ANR21-CE28-0015-01.

## AUTHOR CONTRIBUTIONS

BM and VLM Designed Research; MM and AT Acquired Data; VLM, JG, MM and BLG Analyzed Data; VLM, MM, LL, DP, ALG, LHA and BM Interpreted Results; VLM, BM and LHA Wrote the first draft of the paper; all authors contributed to the manuscript’s final Writing.

## SUPPLEMENTARY MATERIAL

## METHODS

### Audio corpus

We analyzed audio recordings from three corpora of naturalistic sounds: speech, music, and environmental sounds. The speech corpus is composed of verses of the New Testament from 17 different languages (www.faithcomesbyhearing.com). We selected the specific versions of the recordings that did not contain any sound effects but only plain speech (Table S1).

The music corpus comes from the “GTZAN Dataset – Music Genre Classification” (*GTZAN Dataset - Music Genre Classification*, s. f.); “the MNIST of sounds,” a collection of 10 musical genres comprising blues, classical, country, disco, hiphop, jazz, metal, pop, reggae, and rock. Each genre contains 100 audio files, each with a length of 30 s, sampled at 22.05 kHz.

The corpus of environmental sounds comes from the “Making Sense of Sound” classification challenge (Kroos et al., 2019), containing a broad range of sounds from everyday life. This corpus is composed of five categories: nature, human, music, effects, and urban, with 300 sounds in each category. All samples were 5 s long, sampled at 44.1 kHz, and peak-normalized.

The audios of each corpus were converted from multi to single channel and resampled to 16 kHz. Each file was then cut into a 3- s- long segment, and its amplitude envelope was extracted. Amplitude envelope corresponds to the slow overall amplitude fluctuations of the signal over time. It was computed with custom MATLAB scripts developed for speech signals analyses (Gilbert & Lorenzi, 2006). The sound signal was decomposed into 32 narrow frequency bands using a cochlear model, and the absolute value of the Hilbert transform was computed for each of these narrowband signals. The broadband temporal amplitude envelope resulted from the summation of these band-pass–filtered signals and was down sampled to 512Hz.

For 9 languages of the speech corpus, we obtained the precise timing of each phonemes present in the audio files (Table S1). To do so, we first obtained the transcript of each audio file using the speech to text model whisper-large-v3 from OpenAI available through the HugginFace platform (Radford et al., 2022). Following this first step, we used the WebMAUS online tool (Kisler et al., 2017) to force align the sounds and transcripts and obtained for each audio a textgrid file with the timestamps of each phonemes. For languages available in the Mass corpus (Zanon Boito et al., 2020), the textgrid files were directly retrieved without the need for the previous steps.

For each language and each phoneme, the corresponding amplitude envelope time series were segmented into epochs spanning from 1 second before to 1 second after the phoneme of interest. Epochs corresponding to the same phoneme were then grouped together and subjected to time-frequency decomposition. Distinct phonemes were further categorized based on whether they belonged to the vowel or consonant group.

### Participants

Two cohorts of patients with pharmacoresistant epilepsy underwent stereo-electroencephalographic (SEEG) recordings during their presurgical evaluation at Hôpital de la Timone (Marseille, France). The first cohort included 18 patients (10 females), while the second comprised 2 patients (1 female). Unless otherwise stated, all analyses were performed on the first cohort. All patients were French native speakers and the neuropsychological assessments confirmed that they had intact language functions and normal hearing. None of the patients had their epileptogenic zones located in the auditory areas, as determined by experienced epileptologists. The study received approval from the Institutional Review Board of the French Institute of Health (IRB00003888) in accordance with the Declaration of Helsinki. All patients provided written informed consent before the experimental session. Participation was voluntary, and none of the patients were involved in a clinical trial.

### Recordings

Stereo-electroencephalographic (SEEG) recordings SEEG recordings were performed using depth electrodes, implanted stereotactically (Talairach et al., 1992, Alcis, Besançon, France, and Dixi Medical, Chaudefontaine, France). All the patients had one electrode located in the auditory cortex, implanted orthogonally to the cortical surface, targeting the Heschl’s gyrus and the planum temporale (Figure 1a). The first cohort consisted of 13 unilateral implantations (10 left, 3 right) and 5 bilateral implantations, resulting in a total of 23 analyzed electrodes (N=23). The second cohort had N=3 electrodes implanted in the region of interest, including one patient with a bilateral implantation. The diameter of the electrodes was 0.8 mm and they had and between 8 and 18 contacts of 2 mm length and separated from each other by 1.5 mm. The recording reference and ground were chosen by the clinical staff as two consecutive sEEG contacts on the same shaft both located in the white matter and/or at distance from any epileptic activity. Patients were recorded either in an insulated Faraday cage or in the bedroom. In the Faraday cage, they laid comfortably in a chair, the room was sound attenuated, and data were recorded using a 256-channel amplifier (Brain Products), sampled at 1 kHz and high-pass filtered at 0.016 Hz. In the bedroom, data were recorded using a 256-channel Natus amplifier (DeltaMed system), sampled at 512 Hz, and high-pass filtered at 0.16 Hz. To precisely localize the channels, a procedure similar to the one used in the iELVis toolbox and in the FieldTrip toolbox was applied (Groppe et al., 2017; Stolk et al., 2018). First, we manually identified the location of each channel centroid on the post-implant CT scan using the Gardel software (Medina Villalon et al., 2018). Second, we performed volumetric segmentation and cortical reconstruction on the pre-implant MRI with the Freesurfer image analysis suite (documented and freely available for download online at http://surfer.nmr.mgh.harvard.edu/). This segmentation of the pre-implant MRI with SPM12 provides us with both the tissue probability maps (i.e. gray, white, and cerebrospinal fluid [CSF] probabilities) and the indexed-binary representations (i.e. either gray, white, CSF, bone, or soft tissues). This information allowed us to reject electrodes not located in the brain. Third, the post-implant CT scan was coregistered to the pre- implant MRI via a rigid affine transformation and the pre-implant MRI was registered to MNI152 space, via a linear and a non-linear transformation from SPM12 methods (Penny et al., 2007), through the FieldTrip toolbox (Oostenveld et al., 2011). Fourth, applying the corresponding transformations, we mapped channel locations to the pre-implant MRI brain that was labeled using the volume-based Human Brainnetome Atlas (Fan et al., 2016).

### Experimental paradigm

For the first cohort, SEEG activity was recorded during rest and a passive listening task. The resting condition consisted of three three-minute intervals, during which patients remained awake without specific instructions to open or close their eyes. The task consisted on passively listening to a ∼10 minutes-long storytelling (577 s, *La sorcière de la rue Mouffetard*), while recording SEEG activity. The fundamental frequency was measured using Praat’s autocorrelation algorithm (male voice, median F_0_ = 139 Hz). The second cohort of two patients performed the same tasks and additionally listened to a second story (613 s, *Le Petit Prince*) with a higher fundamental frequency (female voice, median F_0_ = 189 Hz).

The experiment was done either in an insulated Faraday cage or in the patient’s bedroom. In the former, a sound Blaster X-Fi Xtreme Audio, an amplifier Yamaha P2040 and Yamaha loudspeakers (NS 10M) were used for sound presentation. In the latter, stimuli were presented using a Sennheiser HD 25 headphone set. Sound stimuli were presented at ∼75 dBA, with 44.1 kHz sample rate and 16 bits resolution.

To analyze the brain activity in response to the speech, we computed the envelope of the audio signal as main feature, following the same approach as for the audio corpus.

### Independent Component Analysis

ICA aims to solve the ‘cocktail party’ problem by separating N statistically independent sources that have been mixed on M recording contacts (Herreras et al., 2022; López-Madrona, Trébuchon, Bénar, et al., 2024; López-Madrona, Trébuchon, Mindruta, et al., 2024; Makarova et al., 2025). It assumes immobility of the neural sources in space, recording session. Each recorded signal *u_m_*(*t*) is thus modeled as the sum of *N* independent i.e., that the contribution of one source to the recording contacts is the same throughout the sources (*s_n_*(*t*)) multiplied by a constant factor (*V_mn_*)

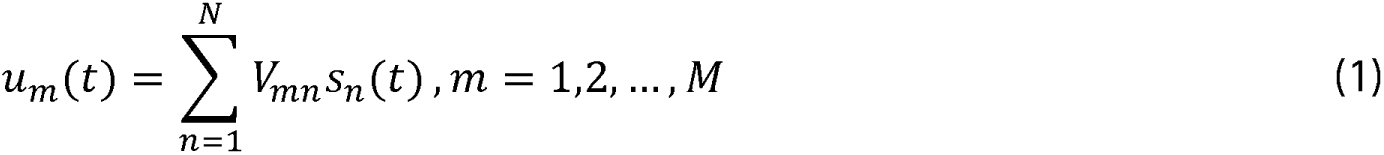

where *u_m_*(*t*) is the SEEG data, *V_mn_* the ICA weights or spatial profile of each source, *M* the number of contacts, *N* the number of sources and *s_n_*(*t*) the obtained independent components (“SEEG-ICs”).

For each electrode, we obtained as many components as available contacts (N=M), without any prior dimension reduction (Artoni et al., 2018). We used EEGLAB (Delorme & Makeig, 2004) to compute ICA using the infomax algorithm (Bell & Sejnowski, 1995). From all the components, we manually rejected all the SEEG-ICs related to the reference or remote sources. They were characterized by a flat spatial profile with a similar contribution to all the contacts of the electrode (López-Madrona, Trébuchon, Bénar, et al., 2024). All the other SEEG-ICs were considered as putative neural sources. All the SEEG-ICs were z-scored.

### Spectral analysis

Power spectra were estimated using the multitaper method (Thomson, 1982), with a frequency resolution of 0.25 Hz. To characterize the power of each source, we applied the *fooof* approach (Donoghue et al., 2020), which separates the periodic and aperiodic (1/f-like) components of the spectra. This method allows for the analysis of oscillatory power independently of 1/f-related changes (Voytek et al., 2015). The aperiodic component was modeled using a Lorentzian function, with offset, knee, and exponent (curvature) as key parameters. Then, *fooof* identified oscillatory peaks exceeding the aperiodic component and fit them individually using Gaussian functions, yielding estimates of power, center frequency, and bandwidth for each detected oscillation.

For the analysis in Figure 2e-g, we applied this approach to identify significant oscillatory peaks, indicating the presence of oscillations in the time course. The *fooof* fit was performed in the 1-50 Hz range, with a minimum peak bandwidth of 0.5 Hz (twice the frequency resolution) and a minimum amplitude threshold of twice the standard deviation of the aperiodic-removed power spectrum. If multiple peaks were detected with overlapping bandwidths, they were merged, retaining the frequency and relative power of the highest peak.

Since the aperiodic activity during auditory stimulation differs from the resting state (López-Madrona, Trébuchon, Bénar, et al., 2024), we removed this component from both conditions using the fooof approach (Donoghue et al., 2020).To account for differences in the 1/f distribution between rest and auditory stimulation (López-Madrona, Trébuchon, Bénar, et al., 2024) (Figure 2l), we recomputed the *fooof* fit separately for two frequency bands: 1-28 Hz and 28-150 Hz (Podvalny et al., 2015). A distinct aperiodic component was fitted for each SEEG-IC and frequency band, which was subsequently removed from the power spectra.

#### Statistical test for individual analysis

Recordings (rest and listening task) were segmented into 10-second windows, and power spectra were computed for each window. The power spectrum for each task was then averaged, and the ratio was calculated as (speech – rest) / rest, where “speech” refers to the listening condition. The p-value for each frequency was obtained using a permutation test (N = 1001 permutations), randomly reassigning “rest” or “listening task” labels while maintaining the same proportion in each iteration. Multiple comparisons were corrected using False Discovery Rate (FDR).

#### Statistical test for group analysis

A single power spectrum was averaged for each SEEG-IC and condition, and the ratio was computed as described above. A t-test was performed at each frequency to determine whether the ratio differed significantly from zero, with FDR correction for multiple comparisons.

### Time-frequency analysis

The time-frequency representation of each envelope was computed using Morlet wavelet transforms with 7 cycles per wavelet. Each frequency band was normalized separately using a z-score computed over the entire duration of the audio recording. Time-frequency maps were then segmented into ‘trials’, time-locked to phoneme onsets, and categorized into two conditions: vowels and consonants. For each language, we averaged the time-frequency responses across phonemes within each condition. To assess differences in 30–50 Hz activity between conditions, we extracted the maximum power in this frequency range following phoneme onset for each language and condition. Group-level statistical comparisons across languages were performed using a paired t-test.

### Granger Causality

To estimate whether a SEEG-IC was related to the speech signal, we computed the Granger Causality (GC) between them (Bressler & Seth, 2011; Granger, 1969; López-Madrona et al., 2019). GC assumes that, if there is a linear relation between *x*(*t*) and *y*(*t*) (e.g., between the speech envelope and the neural activity, respectively), then the past activity of *x* (*x*(*t-τ*)) may predict the dynamics of *y*(*t*). It is based on autoregressive models, where *x*(*t*) and *y*(*t*) are modeled by their own past and a residual. Briefly, if there is a directionality from *x*(*t*) to *y*(*t*) the inclusion of the previous values of *x*(*t*) in the model of *y*(*t*) would reduce the error of the prediction. Then, the GC in the temporal domain from *x*(*t*) to *y*(*t*) would be determined by the F-statistics:

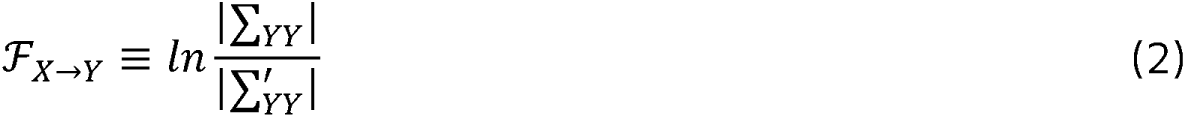

where 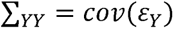 and 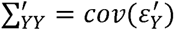 are the residual covariance matrices of the autoregressive models of *y*(*t*) before and after including the past values of *x*(*t*) respectively.

While equation (2) refers to the GC in the temporal domain, it is also possible to decompose it by frequency, estimating a spectral version of the GC (Geweke, 1982). The advantage of the spectral GC is to identify which are the frequencies of *y*(*t*) that can be explained by the same oscillations from *x*(*t*).

We used the MVGC toolbox (Barnett & Seth, 2014) to computed the unconditional GC between the speech envelope and each SEEG-IC independently (i.e., only two signals in a single model). We downsampled the signals to 360 Hz and used a sliding window of 10 seconds, with an overlap of 80% and a model order of 15. The GC associated to each pair of signals was the average across time-windows. The statistical significance was obtained with a surrogate data analysis (n=1001) by shifting the speech time course by a random value and breaking the temporal relationship between them. We computed the GC for each surrogate and them approximated them into a gaussian distribution. The significance threshold was set at the 95th percentile of the distribution (pval = .05).

An SEEG-IC was considered speech-driven if it exhibited significant temporal GC from the envelope (Figure 2i). We then analyzed the spectral content of the GC and categorized SEEG-ICs into three groups (Figure 3a): theta SEEG-ICs, if significance was found only at theta frequencies (2-6 Hz); gamma SEEG-ICs, if significance was limited to low- and/or high-gamma bands (30-50 Hz and 100-150 Hz, respectively); and theta-gamma SEEG-ICs, if significance was present in both the theta range and either low- or high-gamma to the SEEG-IC).

### Phase-Amplitude Coupling

To evaluate the non-linear interactions between theta (2-6 Hz) and gamma (30-150 Hz) oscillations, we computed the phase-amplitude coupling (PAC). The PAC measures whether the changes in amplitude of a fast signal (e.g. gamma activity) occur at specific phases of a slower oscillation (e.g. theta). We used a modified version of the modulation index (MI) (Canolty et al., 2006; Tort et al., 2008) to mitigate the errors induced when estimating the phase of oscillations that are not purely sinusoidal (López-Madrona et al., 2020).

For each frequency considered as reference for the phase, we combined a narrowband filter to detect the zero-crossing points (which correspond to the ascendant and descendant slope of the oscillation, or the phases π/2 and 3π/3, respectively) and a broadband filter to find the trough and the peak (phases 0 and π, respectively; Cole and Voytek, 2019). We labelled this signal as *x_θφ_*(*t*) For each frequency considered as reference for the amplitude we bandpass filtered the signal at the frequency of interest with a bandwidth of 20 Hz (Aru et al., 2015) and computed the envelope using the Hilbert transform. This signal was labelled as *x_γA_*(*t*)

To compute the MI, we divided each cycle of *x_θφ_*(*t*) into N bins (16 in this work). As the phase was defined by 4 points (the trough, the peak and the two zero-crossing points), we split each cycle in its 4 corresponding segments and further divided them in N/4 bins (López-Madrona et al., 2020). Note that as the signal is not sinusoidal, the size of bins may differ within the same cycle. We then computed the mean amplitude of *x_γA_*(*t*) at each bin, calling < *x_γA_* >*_φ_*(*j*) the amplitude at the phase bin *j*. From them, we can calculate the entropy *H*, defined by:

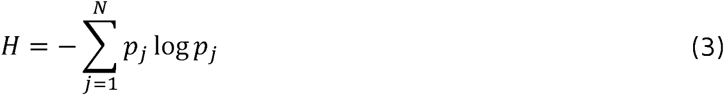

where *N* was set to 16, and *p_j_* is given by

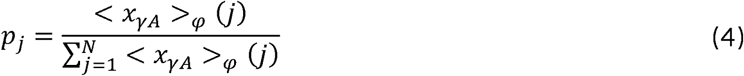

The value of MI was obtained normalizing *H* by the maximum entropy (*H_max_*), given by the uniform distribution *p_j_*=1/*N* (i.e. *H_max_* = log *N*):

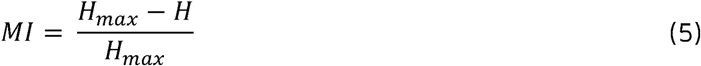

where a MI close to 0 indicates lack of PAC and larger values reflect higher coupling between both signals.

The PAC can be computed between any pair of time-courses, either using the same signal as reference for the phase and amplitude (e.g. PAC within the speech envelope or within an SEEG-IC), or using different signals (e.g. coupling between the phase of the speech envelope and gamma amplitudes of a SEEG-IC, or inter-source PAC between different SEEG-ICs). For each pair of analyzed signals, a comodulogram was obtained as the MI map between certain ranges of frequencies for the phase and amplitude references. The shifting *x_γA_*(*t*) with respect to *x_θφ_*(*t*). This breaks the temporal alignment between both statistical significance was assessed using a surrogate analysis (n=1001 surrogates) by oscillations and the MI obtained can be considered “by chance”. To correct the comodulogram for multiple comparisons, we used a cluster-based approach (Cohen, 2014).

To obtain a single PAC value for low- and high-gamma per SEEG-IC, we selected the maximum comodulogram value within the 2-6 Hz range for the phase and the 30-50 Hz range for the amplitude of low-gamma, or the 100-150 Hz range for high-gamma.

### Cross-Frequency Directionality

While the PAC gives an estimation of the degree of coupling, is it possible to measure the directionality between the slow signal and the amplitude of the fast one (Arinyo-i-Prats et al., 2024; Dupré la Tour et al., 2017; Jiang et al., 2015). We used the cross-frequency directionality (CFD) index (Jiang et al., 2015) to infer whether one of the signals was significantly preceding the other. The CFD, based on the phase-slope index (PSI; Nolte et al., 2008), assumes that if the oscillation of one signal *x*(*t*) at a certain frequency is driving a second one *y*(*t*) with a time delay, then the phase difference between them at that specific delay will change consistently with the frequency of the signals.

Calling *x*(*t*) to the original signal, 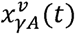 to the power envelope of the signal at a *v* gamma frequency, and being X and 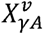 their Fourier transform, respectively, the CFD can be defined by:

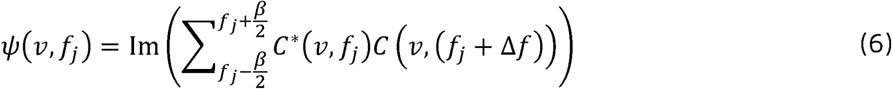

where

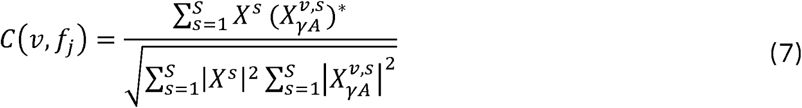

is the complex coherency, *f_j_* is the theta frequency under study, *S* is the number of slope is measured, and it has been fixed at 2 Hz, 4 times the resolution (Δ*f* = 0.5 Hz). segments in which the signal has been divided and *β* is the bandwidth for which the phase

This methodology overcomes some limitations of classical approaches as GC, were the strong differences in signal-to-noise ratio between both components may strongly bias the results (Jiang et al., 2015; Nolte et al., 2010).

We computed the CFD using four different combination of signals: (i) between the phase of the speech envelope and the amplitude of gamma and theta-gamma SEEG-ICs (see Granger Causality section), (ii) between the phase of theta and theta-gamma SEEG-ICs and the amplitude of the speech envelope, (iii) using the phase and amplitude of the same theta-gamma SEEG-IC (intra-source CFD), and (iv) between the phase and amplitude of different SEEG-IC within the same patient (inter-source CFD). To avoid some patients with multiple speech-driven SEEG-ICs bias the results for intra-source and inter-source CFD, we averaged the comodulograms within each electrode, resulting in one single CFD map per electrode.

The statistical significance of the CFD was computed following a pixel-based surrogate approach (Cohen, 2014). In this case, for each surrogated comodulogram (n=1001 surrogates) we kept both the maximal and minimal values of CFD, as the results can be either positive or negative. Thus, we fitted two gaussian distributions and computed both significance thresholds (one for the positive and the other for the negative values) as their 97.5th percentile.

To obtain a single CFD value for low- and high-gamma per SEEG-IC, we selected the absolute maximum value within the 2-6 Hz range for the phase and the 30-50 Hz range for the amplitude of low-gamma, or the 100-150 Hz range for high-gamma.

### Delay speech-brain

The delay between the auditory stimulus and the brain response was estimated as the lag of the cross-correlation between the speech envelope and each SEEG-IC. Separate delay values were computed for theta, low-gamma, and high-gamma by considering only SEEG-ICs with significant GC from the speech envelope at the respective frequency bands. For theta SEEG-ICs, signals were bandpass filtered between 2-6 Hz. For low- and high-gamma SEEG-ICs, signals were filtered at 30-50 Hz and 100-150 Hz, respectively, and their envelopes were extracted using the Hilbert transform (Xu et al., 2023). Both the speech envelope and the filtered SEEG-IC signals were segmented into 10-second windows with a 5-second overlap. Cross-correlation was computed within a maximum lag of 500 ms, and the correlation values were averaged across windows. The maximum correlation value and its corresponding lag were retained.

To assess significance, a surrogate approach was applied (n = 1001 surrogates) by randomly shifting the brain signal relative to the speech envelope before segmentation and repeating the analysis. Only the maximum correlation value from each iteration was retained, regardless of lag. A delay was considered reliable if the original maximum correlation exceeded 95% of the surrogate values (p < .05), and its corresponding lag was taken as the estimated neural response delay to the speech signal.

## Data and code availability

The conditions of our ethics approval do not permit public archiving of anonymized study data. Readers seeking access to the data should contact the lead author. Access will be granted to named individuals in accordance with ethical procedures governing the reuse of clinical data, including completion of a formal data sharing agreement.

All the code used in this work is available at https://github.com/jeremygrd/Phase-Amplitude-Coupling-PAC-speech-Amplitude-Envelope/tree/main (acoustic analysis) and https://github.com/VictorLopezMadrona/Matlab_analysis_neuroscience (brain analysis).

---

**Supplementary Table 1:**
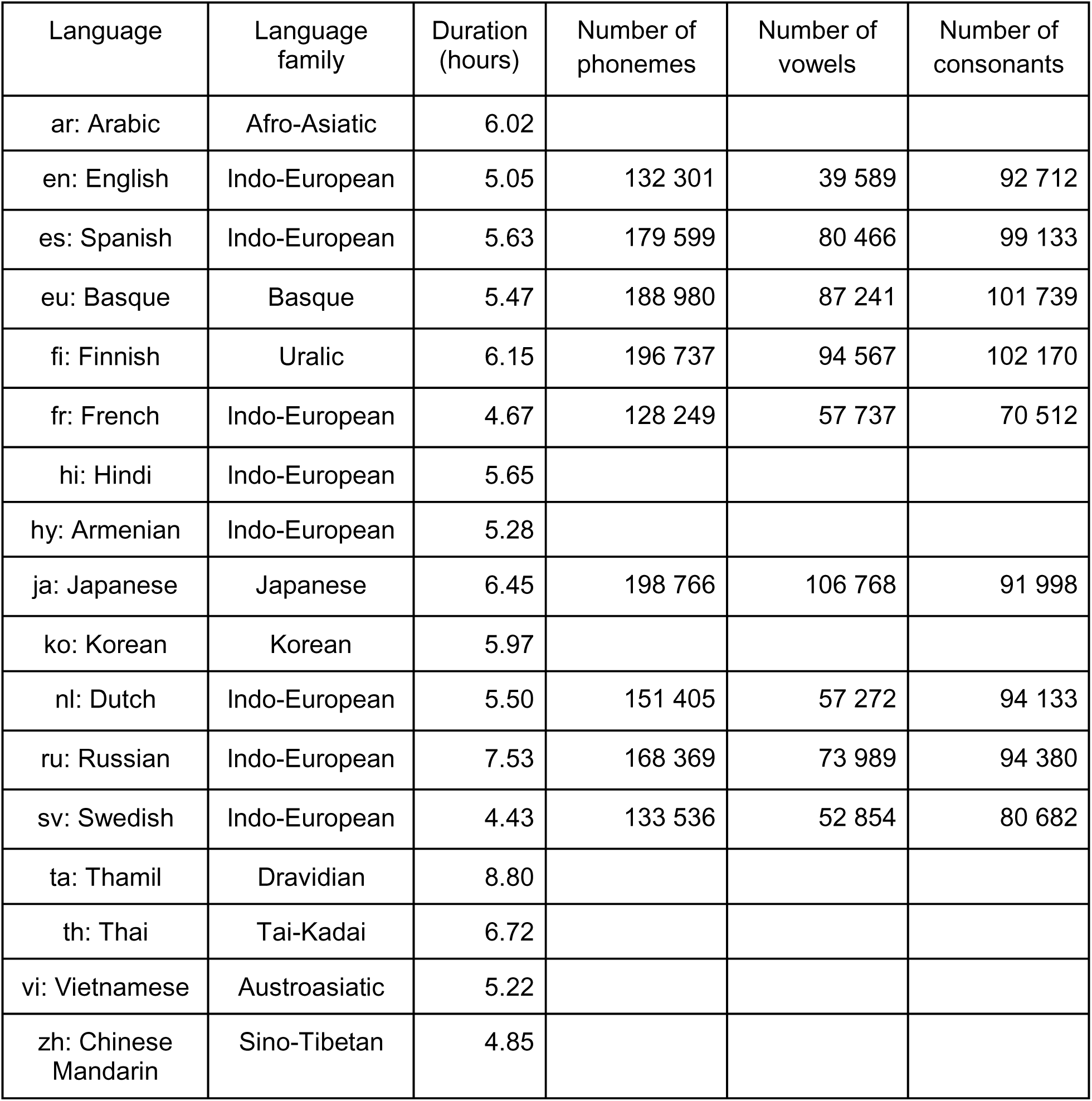
Description of the speech corpus of 17 languages.

| Language | Language family | Duration (hours) | Number of phonemes | Number of vowels | Number of consonants |
| --- | --- | --- | --- | --- | --- |
| ar: Arabic | Afro-Asiatic | 6.02 |  |  |  |
| en: English | Indo-European | 5.05 | 132 301 | 39 589 | 92 712 |
| es: Spanish | Indo-European | 5.63 | 179 599 | 80 466 | 99 133 |
| eu: Basque | Basque | 5.47 | 188 980 | 87 241 | 101 739 |
| fi: Finnish | Uralic | 6.15 | 196 737 | 94 567 | 102 170 |
| fr: French | Indo-European | 4.67 | 128 249 | 57 737 | 70 512 |
| hi: Hindi | Indo-European | 5.65 |  |  |  |
| hy: Armenian | Indo-European | 5.28 |  |  |  |
| ja: Japanese | Japanese | 6.45 | 198 766 | 106 768 | 91 998 |
| ko: Korean | Korean | 5.97 |  |  |  |
| nl: Dutch | Indo-European | 5.50 | 151 405 | 57 272 | 94 133 |
| ru: Russian | Indo-European | 7.53 | 168 369 | 73 989 | 94 380 |
| sv: Swedish | Indo-European | 4.43 | 133 536 | 52 854 | 80 682 |
| ta: Thamil | Dravidian | 8.80 |  |  |  |
| th: Thai | Tai-Kadai | 6.72 |  |  |  |
| vi: Vietnamese | Austroasiatic | 5.22 |  |  |  |
| zh: Chinese Mandarin | Sino-Tibetan | 4.85 |  |  |  |

**Supplementary Table 2:** Anatomical location of the SEEG-ICs. STG: Superior Temporal Gyrus; A22c: caudal area 22; A22r: rostral area 22; STS: Superior Temporal Sulcus; rpSTS: rostroposterior STS. Brainnetome atlas.

| Region | # Theta<br>SEEG-ICs | # Gamma<br>SEEG-ICs | # Theta-Gamma<br>SEEG-ICs |
| --- | --- | --- | --- |
| STG (A41/A42) | 5 | 10 | 7 |
| STG (A22c) | 1 | 2 | 0 |
| STG (A22r) | 1 | 1 | 1 |
| STG (TE1.0 and TE1.2) | 1 | 2 | 3 |
| STS (rpSTS) | 1 | 4 | 3 |
| Insular gyrus | 1 | 0 | 5 |
| Other | 0 | 7 | 5 |
| **Total:** | **10** | **26** | **24** |

**Supplementary Figure 1:**
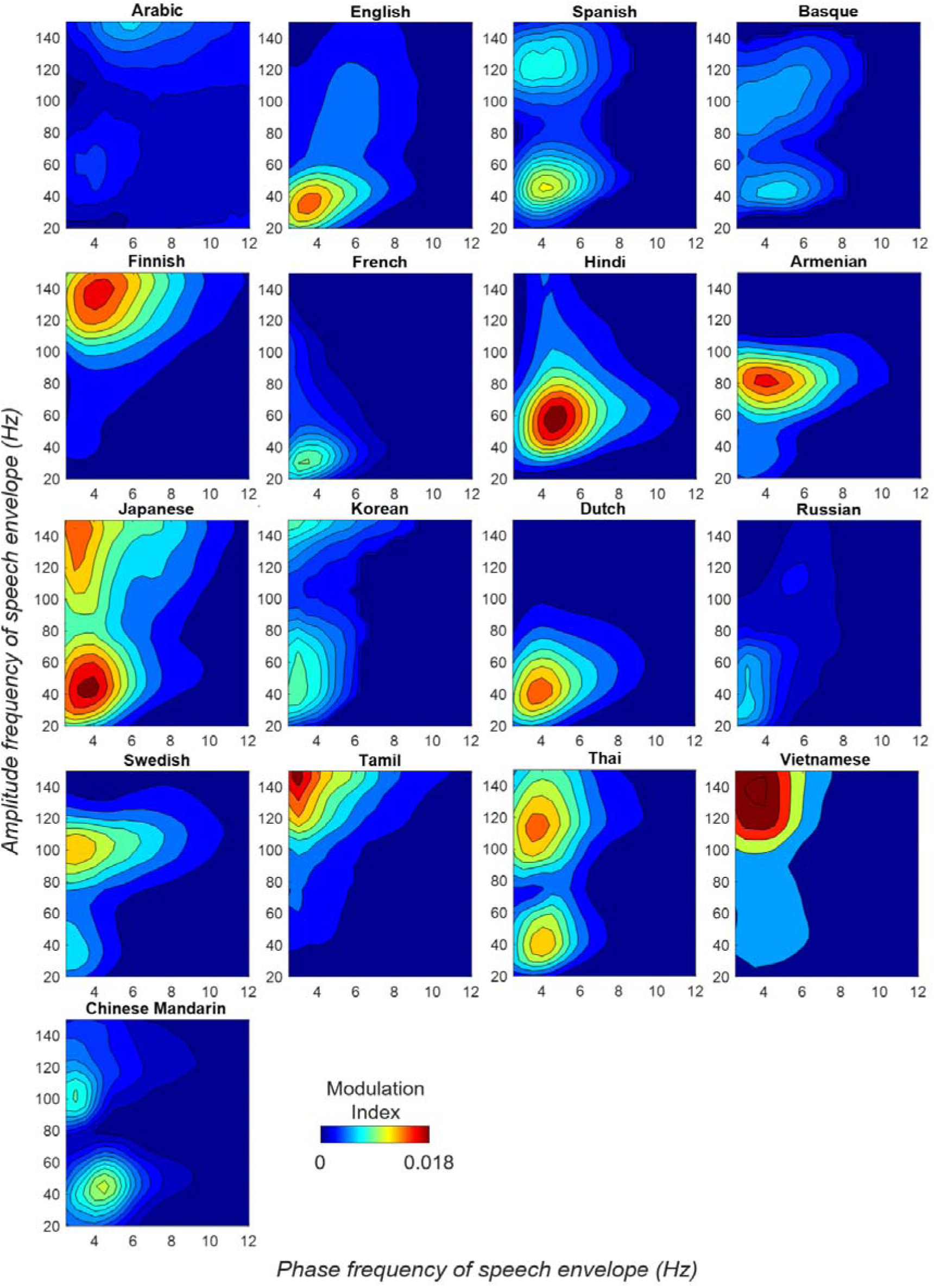
PAC of the speech envelope across 17 languages. PAC comodulogram between the phase and the amplitude of the speech envelope of naturalistic, discourse-level speech (p<.05, surrogate tests).

**Supplementary Figure 2:**
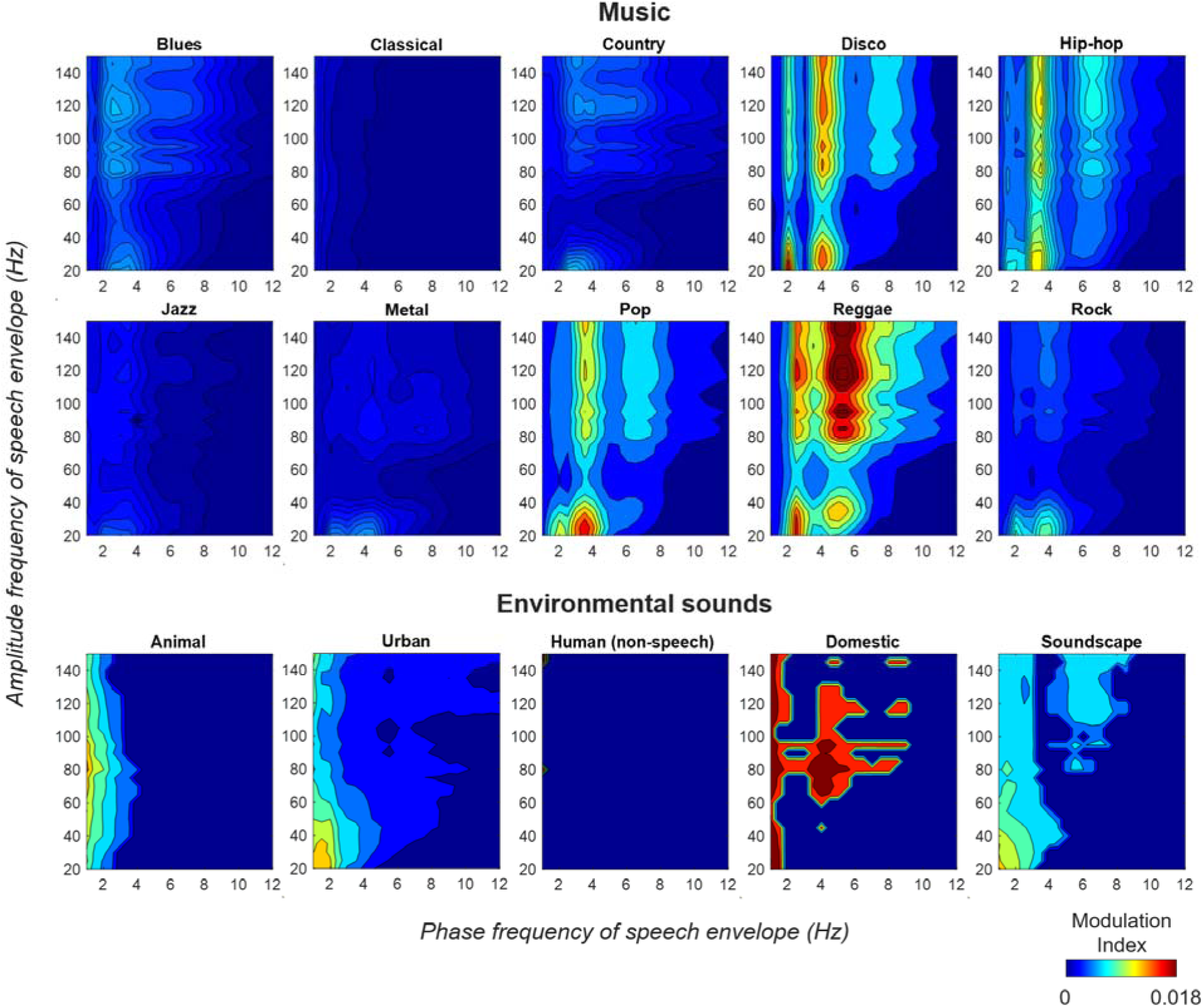
PAC of the envelope of various naturalistic musical and environmental sounds. PAC comodulogram between the phase and the amplitude of the envelope of (top) 10 musical genres and (bottom) 5 categories of environmental sounds.

**Supplementary Figure 3:**
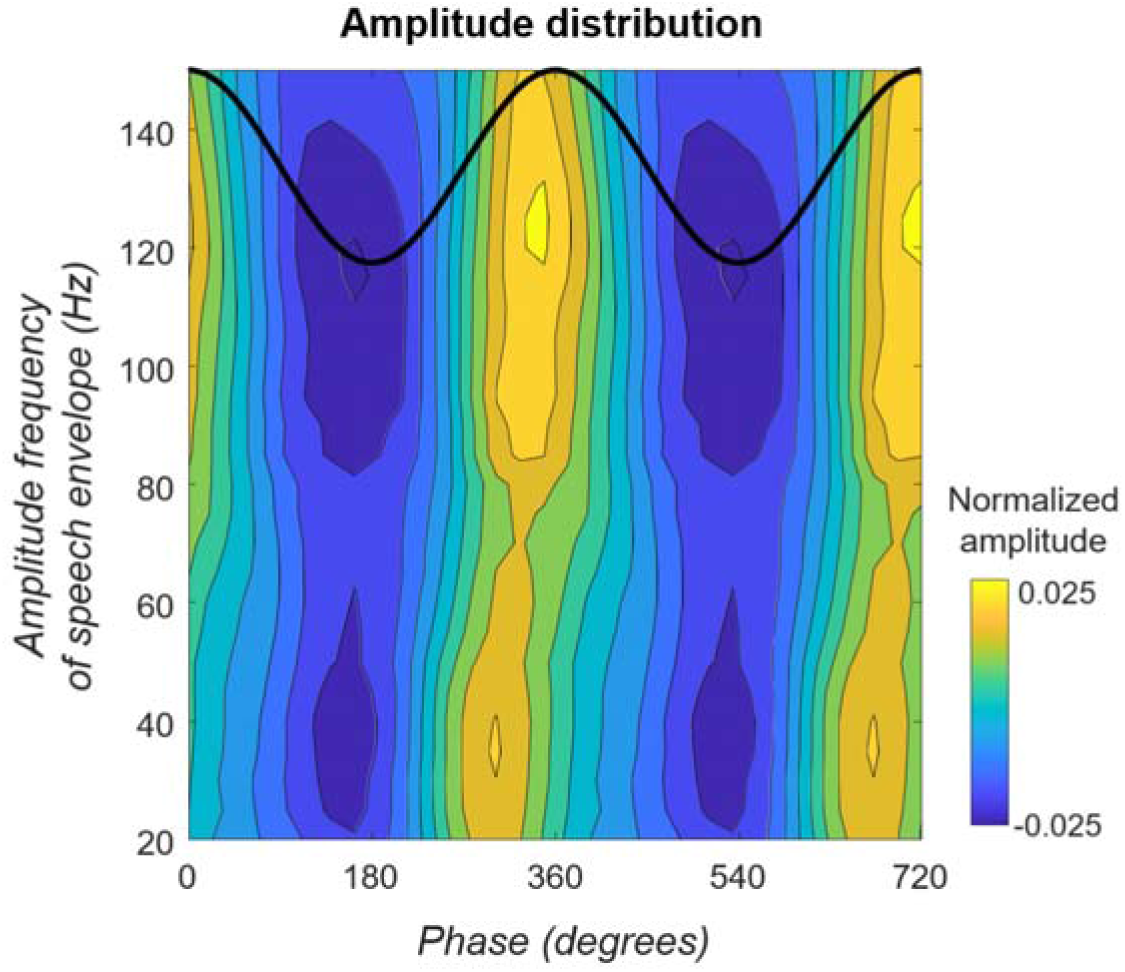
Preferred phase of PAC. Averaged normalized amplitude distribution across two 4 Hz cycles of the speech envelope of the french male voice. The black line represents a reference for the phase signal.

**Supplementary Figure 4:**
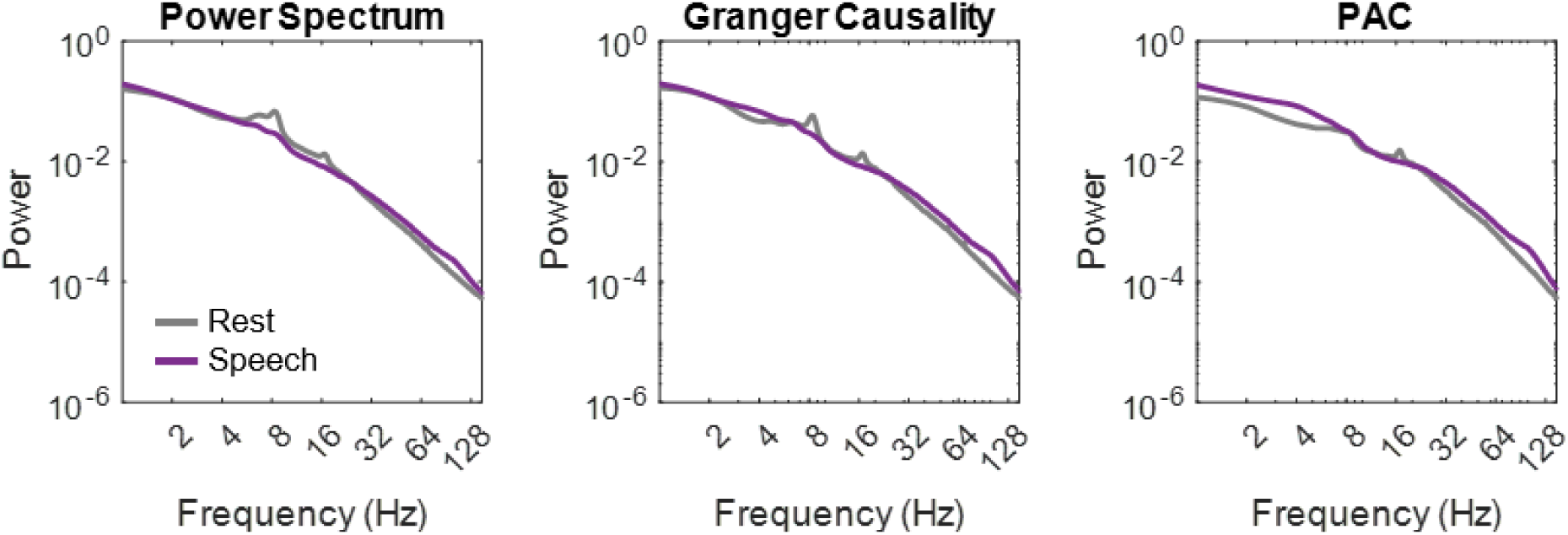
Non-normalized power spectrum in rest and speech. Power spectra during rest and speech for all components with: (left) significant changes in power between conditions; (middle) significant GC from the speech envelope; (right) significant phase-amplitude coupling (PAC) between the phase of theta and the gamma amplitude.

**Supplementary Figure 5.**
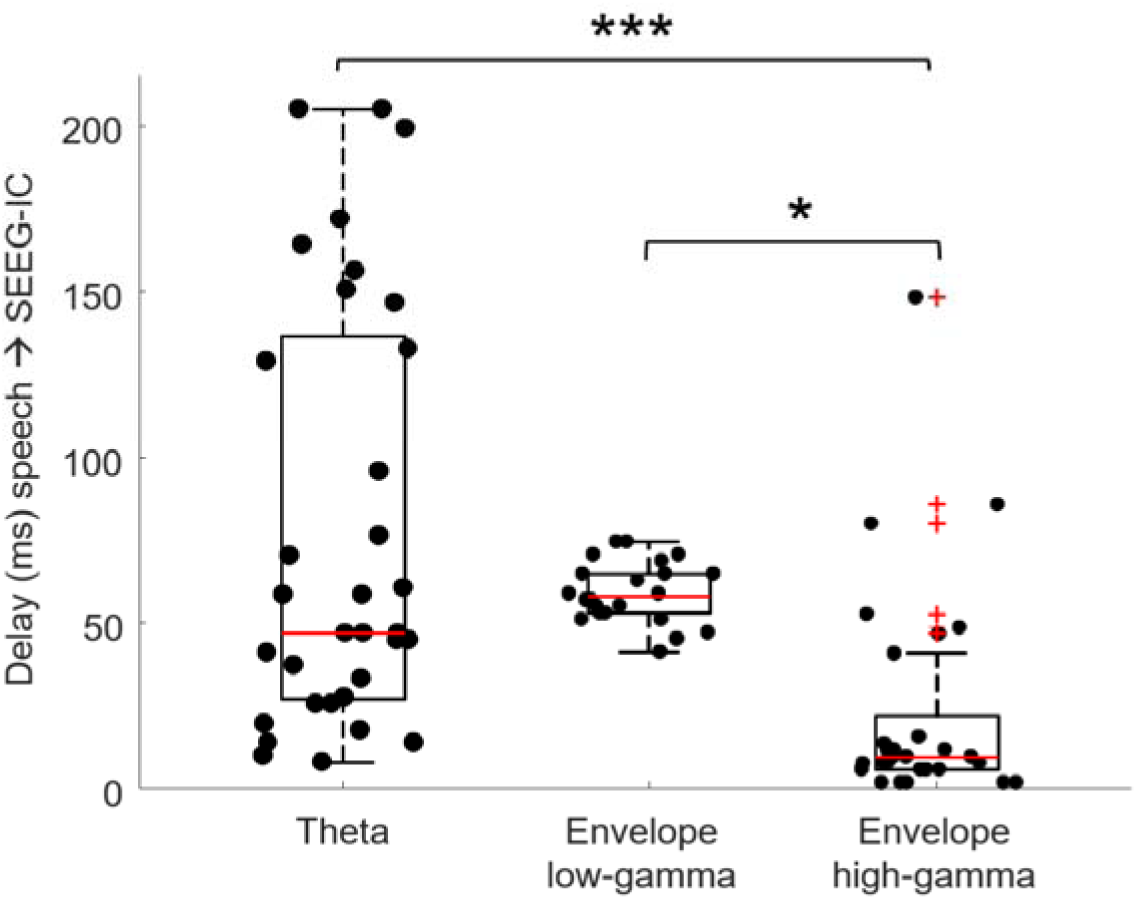
Temporal lag between the speech envelope and auditory activity. Delay estimated using cross-correlations between the speech envelope and speech-driven SEEG-ICs with spectral GC at theta (N=33 SEEG-ICs showing a significant cross-correlation with the speech envelope, see Methods), low-gamma (N=22), or high-gamma (N=29) frequencies. (*/*** p < .05/.001, Tukey’s HSD post-hoc test). Points represent individual SEEG-ICs. Box plots show the median (center line), interquartile range (box), and full data range excluding outliers (whiskers); outliers are marked with red crosses.

## BIBLIOGRAPHY

Ahissar, E., Nagarajan, S., Ahissar, M., Protopapas, A., Mahncke, H., & Merzenich, M. M. (2001). Speech comprehension is correlated with temporal response patterns recorded from auditory cortex. Proceedings of the National Academy of Sciences, 98(23), 13367–13372. 10.1073/pnas.201400998

Akam, T. E., & Kullmann, D. M. (2012). Efficient “Communication through Coherence” Requires Oscillations Structured to Minimize Interference between Signals. PLOS Computational Biology, 8(11), e1002760. 10.1371/journal.pcbi.1002760

Armonaite, K., Bertoli, M., Paulon, L., Gianni, E., Balsi, M., Conti, L., & Tecchio, F. (2022). Neuronal Electrical Ongoing Activity as Cortical Areas Signature: An Insight from MNI Intracerebral Recording Atlas. Cerebral Cortex, 32(13), 2895–2906. 10.1093/cercor/bhab389

Arnal, L. H., Kleinschmidt, A., Spinelli, L., Giraud, A.-L., & Mégevand, P. (2019). The rough sound of salience enhances aversion through neural synchronisation. Nature Communications, 10(1), 3671. 10.1038/s41467-019-11626-7

Atanasova, T., Gross, J., Rimmele, J., & Keitel, A. (2025). The involvement of endogenous brain rhythms in speech processing. OSF. 10.31234/osf.io/rukwp_v1

Attaheri, A., Choisdealbha, Á. N., Di Liberto, G. M., Rocha, S., Brusini, P., Mead, N., Olawole-Scott, H., Boutris, P., Gibbon, S., Williams, I., Grey, C., Flanagan, S., & Goswami, U. (2022). Delta- and theta-band cortical tracking and phase-amplitude coupling to sung speech by infants. NeuroImage, 247, 118698. 10.1016/j.neuroimage.2021.118698

Baillet, S. (2017). Magnetoencephalography for brain electrophysiology and imaging. Nature Neuroscience, 20(3), Article 3. 10.1038/nn.4504

Baroni, F., Morillon, B., Trébuchon, A., Liégeois-Chauvel, C., Olasagasti, I., & Giraud, A.-L. (2020). Converging intracortical signatures of two separated processing timescales in human early auditory cortex. NeuroImage, 218, 116882. 10.1016/j.neuroimage.2020.116882

Bidelman, G. M. (2018). Subcortical sources dominate the neuroelectric auditory frequency-following response to speech. NeuroImage, 175, 56–69. 10.1016/j.neuroimage.2018.03.060

Bragin, A., Jandó, G., Nádasdy, Z., Hetke, J., Wise, K., & Buzsáki, G. (1995). Gamma (40-100 Hz) oscillation in the hippocampus of the behaving rat. The Journal of Neuroscience: The Official Journal of the Society for Neuroscience, 15(1 Pt 1), 47–60.

Brugge, J. F., Nourski, K. V., Oya, H., Reale, R. A., Kawasaki, H., Steinschneider, M., & Matthew A. Howard, I. I. I. (2009). Coding of Repetitive Transients by Auditory Cortex on Heschl’s Gyrus. Journal of Neurophysiology. 10.1152/jn.91346.2008

Buzsáki, G., Anastassiou, C. A., & Koch, C. (2012). The origin of extracellular fields and currents—EEG, ECoG, LFP and spikes. Nature Reviews Neuroscience, 13(6), 407–420. 10.1038/nrn3241

Buzsáki, G., & Draguhn, A. (2004). Neuronal Oscillations in Cortical Networks. Science, 304(5679), 1926–1929. 10.1126/science.1099745

Canolty, R. T., Edwards, E., Dalal, S. S., Soltani, M., Nagarajan, S. S., Kirsch, H. E., Berger, M. S., Barbaro, N. M., & Knight, R. T. (2006). High Gamma Power Is Phase-Locked to Theta Oscillations in Human Neocortex. Science (New York, N.Y.), 313(5793), 1626–1628. 10.1126/science.1128115

Canolty, R. T., & Knight, R. T. (2010). The functional role of cross-frequency coupling. Trends in Cognitive Sciences, 14(11), 506–515. 10.1016/j.tics.2010.09.001

Capilla, A., Arana, L., García-Huéscar, M., Melcón, M., Gross, J., & Campo, P. (2022). The natural frequencies of the resting human brain: An MEG-based atlas. NeuroImage, 258, 119373. 10.1016/j.neuroimage.2022.119373

Chait, M., Greenberg, S., Arai, T., Simon, J. Z., & Poeppel, D. (2015). Multi-time resolution analysis of speech: Evidence from psychophysics. Frontiers in Neuroscience, 9. 10.3389/fnins.2015.00214

Chalas, N., Meyer, L., Lo, C.-W., Park, H., Kluger, D. S., Abbasi, O., Kayser, C., Nitsch, R., & Gross, J. (2024). Dissociating prosodic from syntactic delta activity during natural speech comprehension. Current Biology, 34(15), 3537–3549.e5. 10.1016/j.cub.2024.06.072

Chalkiadakis, D., Sánchez-Claros, J., López-Madrona, V. J., Canals, S., & Mirasso, C. R. (2025). The role of feedforward and feedback inhibition in modulating theta-gamma cross-frequency interactions in neural circuits. PLOS Computational Biology, 21(8), e1013363. 10.1371/journal.pcbi.1013363

Chang, A., Li, Y., Roman, I. R., & Poeppel, D. (2025). *Spectrotemporal Modulation: Efficient and Interpretable Feature Representation for Classifying Speech, Music, and Environmental Sounds* (arXiv:2505.23509). arXiv. 10.48550/arXiv.2505.23509

Coffey, E. B. J., Nicol, T., White-Schwoch, T., Chandrasekaran, B., Krizman, J., Skoe, E., Zatorre, R. J., & Kraus, N. (2019). Evolving perspectives on the sources of the frequency-following response. Nature Communications, 10(1), 5036. 10.1038/s41467-019-13003-w

Colgin, L. L., Denninger, T., Fyhn, M., Hafting, T., Bonnevie, T., Jensen, O., Moser, M.-B., & Moser, E. I. (2009). Frequency of gamma oscillations routes flow of information in the hippocampus. Nature, 462(7271), 353–357. 10.1038/nature08573

Coupé, C., Oh, Y. M., Dediu, D., & Pellegrino, F. (2019). Different languages, similar encoding efficiency: Comparable information rates across the human communicative niche. Science Advances, 5(9), eaaw2594. 10.1126/sciadv.aaw2594

Di Liberto, G. M., O’Sullivan, J. A., & Lalor, E. C. (2015). Low-Frequency Cortical Entrainment to Speech Reflects Phoneme-Level Processing. Current Biology, 25(19), 2457–2465. 10.1016/j.cub.2015.08.030

Ding, N., Patel, A. D., Chen, L., Butler, H., Luo, C., & Poeppel, D. (2017). Temporal modulations in speech and music. Neuroscience & Biobehavioral Reviews, 81, 181–187. 10.1016/j.neubiorev.2017.02.011

Ding, N., & Simon, J. Z. (2014). Cortical entrainment to continuous speech: Functional roles and interpretations. Frontiers in Human Neuroscience, 8. 10.3389/fnhum.2014.00311

Doelling, K. B., Arnal, L. H., Ghitza, O., & Poeppel, D. (2014). Acoustic landmarks drive delta–theta oscillations to enable speech comprehension by facilitating perceptual parsing. NeuroImage, 85, 761–768. 10.1016/j.neuroimage.2013.06.035

Dogonasheva, O., Zakharov, D., Giraud, A.-L., & Gutkin, B. (2025). *Neuro-oscillatory models of cortical speech processing* (arXiv:2502.12935). arXiv. 10.48550/arXiv.2502.12935

Donoghue, T., Haller, M., Peterson, E. J., Varma, P., Sebastian, P., Gao, R., Noto, T., Lara, A. H., Wallis, J. D., Knight, R. T., Shestyuk, A., & Voytek, B. (2020). Parameterizing neural power spectra into periodic and aperiodic components. Nature Neuroscience, 23(12), Article 12. 10.1038/s41593-020-00744-x

Etard, O., & Reichenbach, T. (2019). Neural Speech Tracking in the Theta and in the Delta Frequency Band Differentially Encode Clarity and Comprehension of Speech in Noise. Journal of Neuroscience, 39(29), 5750–5759. 10.1523/JNEUROSCI.1828-18.2019

Fan, L., Li, H., Zhuo, J., Zhang, Y., Wang, J., Chen, L., Yang, Z., Chu, C., Xie, S., Laird, A. R., Fox, P. T., Eickhoff, S. B., Yu, C., & Jiang, T. (2016). The Human Brainnetome Atlas: A New Brain Atlas Based on Connectional Architecture. Cerebral Cortex (New York, NY), 26(8), 3508–3526. 10.1093/cercor/bhw157

Fitch, W. T. (2000). The evolution of speech: A comparative review. Trends in Cognitive Sciences, 4(7), 258–267. 10.1016/S1364-6613(00)01494-7

Fontolan, L., Morillon, B., Liegeois-Chauvel, C., & Giraud, A.-L. (2014). The contribution of frequency-specific activity to hierarchical information processing in the human auditory cortex. Nature Communications, 5(1), Article 1. 10.1038/ncomms5694

Forseth, K. J., Hickok, G., Rollo, P. S., & Tandon, N. (2020). Language prediction mechanisms in human auditory cortex. Nature Communications, 11(1), 5240. 10.1038/s41467-020-19010-6

Frauscher, B., von Ellenrieder, N., Zelmann, R., Doležalová, I., Minotti, L., Olivier, A., Hall, J., Hoffmann, D., Nguyen, D. K., Kahane, P., Dubeau, F., & Gotman, J. (2018). Atlas of the normal intracranial electroencephalogram: Neurophysiological awake activity in different cortical areas. Brain: A Journal of Neurology, 141(4), 1130–1144. 10.1093/brain/awy035

Freeman, W. J., & Rogers, L. J. (2002). Fine Temporal Resolution of Analytic Phase Reveals Episodic Synchronization by State Transitions in Gamma EEGs. Journal of Neurophysiology, 87(2), 937–945. 10.1152/jn.00254.2001

Galambos, R., Makeig, S., & Talmachoff, P. J. (1981). A 40-Hz auditory potential recorded from the human scalp. Proceedings of the National Academy of Sciences, 78(4), 2643–2647. 10.1073/pnas.78.4.2643

Ghitza, O. (2011). Linking Speech Perception and Neurophysiology: Speech Decoding Guided by Cascaded Oscillators Locked to the Input Rhythm. Frontiers in Psychology, 2. 10.3389/fpsyg.2011.00130

Ghitza, O. (2013). The theta-syllable: A unit of speech information defined by cortical function. Frontiers in Psychology, 4. 10.3389/fpsyg.2013.00138

Ghitza, O. (2014). Behavioral evidence for the role of cortical θ oscillations in determining auditory channel capacity for speech. Frontiers in Psychology, 5, 652. 10.3389/fpsyg.2014.00652

Ghitza, O. (2017). Acoustic-driven delta rhythms as prosodic markers. Language, Cognition and Neuroscience, 32(5), 545–561. 10.1080/23273798.2016.1232419

Giraud, A.-L., Kleinschmidt, A., Poeppel, D., Lund, T. E., Frackowiak, R. S. J., & Laufs, H. (2007). Endogenous Cortical Rhythms Determine Cerebral Specialization for Speech Perception and Production. Neuron, 56(6), 1127–1134. 10.1016/j.neuron.2007.09.038

Giraud, A.-L., & Poeppel, D. (2012). Cortical oscillations and speech processing: Emerging computational principles and operations. Nature Neuroscience, 15(4), Article 4. 10.1038/nn.3063

Giroud, J., Lerousseau, J. P., Pellegrino, F., & Morillon, B. (2023). The channel capacity of multilevel linguistic features constrains speech comprehension. Cognition, 232, 105345. 10.1016/j.cognition.2022.105345

Giroud, J., Trébuchon, A., Mercier, M., Davis, M. H., & Morillon, B. (2024). The human auditory cortex concurrently tracks syllabic and phonemic timescales via acoustic spectral flux. Science Advances, 10(51), eado8915. 10.1126/sciadv.ado8915

Giroud, J., Trébuchon, A., Schön, D., Marquis, P., Liegeois-Chauvel, C., Poeppel, D., & Morillon, B. (2020). Asymmetric sampling in human auditory cortex reveals spectral processing hierarchy. PLoS Biology, 18(3), e3000207. 10.1371/journal.pbio.3000207

Gourévitch, B., Martin, C., Postal, O., & Eggermont, J. J. (2020). Oscillations in the auditory system and their possible role. Neuroscience & Biobehavioral Reviews, 113, 507–528. 10.1016/j.neubiorev.2020.03.030

Greenberg, S., Carvey, H., Hitchcock, L., & Chang, S. (2003). Temporal properties of spontaneous speech—A syllable-centric perspective. Journal of Phonetics, 31(3), 465–485. 10.1016/j.wocn.2003.09.005

Groppe, D. M., Bickel, S., Keller, C. J., Jain, S. K., Hwang, S. T., Harden, C., & Mehta, A. D. (2013). Dominant frequencies of resting human brain activity as measured by the electrocorticogram. NeuroImage, 79, 223–233. 10.1016/j.neuroimage.2013.04.044

Gross, J., Hoogenboom, N., Thut, G., Schyns, P., Panzeri, S., Belin, P., & Garrod, S. (2013). Speech Rhythms and Multiplexed Oscillatory Sensory Coding in the Human Brain. PLOS Biology, 11(12), e1001752. 10.1371/journal.pbio.1001752

Hamilton, L. S., Edwards, E., & Chang, E. F. (2018). A Spatial Map of Onset and Sustained Responses to Speech in the Human Superior Temporal Gyrus. Current Biology: CB, 28(12), 1860–1871.e4. 10.1016/j.cub.2018.04.033

Herreras, O., Makarova, J., & Makarov, V. A. (2015). New uses of LFPs: Pathway-specific threads obtained through spatial discrimination. Neuroscience, 310, 486–503. 10.1016/j.neuroscience.2015.09.054

Hillebrand, A., Barnes, G. R., Bosboom, J. L., Berendse, H. W., & Stam, C. J. (2012). Frequency-dependent functional connectivity within resting-state networks: An atlas-based MEG beamformer solution. NeuroImage, 59(4), 3909–3921. 10.1016/j.neuroimage.2011.11.005

Hillebrand, A., Tewarie, P., van Dellen, E., Yu, M., Carbo, E. W. S., Douw, L., Gouw, A. A., van Straaten, E. C. W., & Stam, C. J. (2016). Direction of information flow in large-scale resting-state networks is frequency-dependent. Proceedings of the National Academy of Sciences, 113(14), 3867–3872. 10.1073/pnas.1515657113

Hovsepyan, S., Olasagasti, I., & Giraud, A.-L. (2020). Combining predictive coding and neural oscillations enables online syllable recognition in natural speech. Nature Communications, 11(1), 3117. 10.1038/s41467-020-16956-5

Hovsepyan, S., Olasagasti, I., & Giraud, A.-L. (2023). Rhythmic modulation of prediction errors: A top-down gating role for the beta-range in speech processing. PLOS Computational Biology, 19(11), e1011595. 10.1371/journal.pcbi.1011595

Howard, M. F., & Poeppel, D. (2010). Discrimination of speech stimuli based on neuronal response phase patterns depends on acoustics but not comprehension. Journal of Neurophysiology, 104(5), 2500–2511. 10.1152/jn.00251.2010

Howard, M. F., & Poeppel, D. (2012). The neuromagnetic response to spoken sentences: Co-modulation of theta band amplitude and phase. NeuroImage, 60(4), 2118–2127. 10.1016/j.neuroimage.2012.02.028

Hutcheon, B., & Yarom, Y. (2000). Resonance, oscillation and the intrinsic frequency preferences of neurons. Trends in Neurosciences, 23(5), 216–222. 10.1016/S0166-2236(00)01547-2

Hyafil, A., Fontolan, L., Kabdebon, C., Gutkin, B., & Giraud, A.-L. (2015). Speech encoding by coupled cortical theta and gamma oscillations. eLife, 4, e06213. 10.7554/eLife.06213

Hyafil, A., Giraud, A.-L., Fontolan, L., & Gutkin, B. (2015). Neural Cross-Frequency Coupling: Connecting Architectures, Mechanisms, and Functions. Trends in Neurosciences, 38(11), 725–740. 10.1016/j.tins.2015.09.001

Jiang, H., Bahramisharif, A., van Gerven, M. A. J., & Jensen, O. (2015). Measuring directionality between neuronal oscillations of different frequencies. NeuroImage, 118, 359–367. 10.1016/j.neuroimage.2015.05.044

Jiang, H., Bahramisharif, A., van Gerven, M. A. J., & Jensen, O. (2020). Distinct directional couplings between slow and fast gamma power to the phase of theta oscillations in the rat hippocampus. European Journal of Neuroscience, 51(10), 2070–2081. 10.1111/ejn.14644

Kalamangalam, G. P., Long, S., & Chelaru, M. I. (2020). A neurophysiological brain map: Spectral parameterization of the human intracranial electroencephalogram. Clinical Neurophysiology, 131(3), 665–675. 10.1016/j.clinph.2019.11.061

Keitel, A., & Gross, J. (2016). Individual Human Brain Areas Can Be Identified from Their Characteristic Spectral Activation Fingerprints. PLOS Biology, 14(6), e1002498. 10.1371/journal.pbio.1002498

Keitel, A., Gross, J., & Kayser, C. (2018). Perceptually relevant speech tracking in auditory and motor cortex reflects distinct linguistic features. PLOS Biology, 16(3), e2004473. 10.1371/journal.pbio.2004473

Keitel, A., Ince, R. A. A., Gross, J., & Kayser, C. (2017). Auditory cortical delta-entrainment interacts with oscillatory power in multiple fronto-parietal networks. NeuroImage, 147, 32–42. 10.1016/j.neuroimage.2016.11.062

Kendall, T. (2013). Speech Rate, Pause and Sociolinguistic Variation. Palgrave Macmillan UK. 10.1057/9781137291448

Keshishian, M., Akkol, S., Herrero, J., Bickel, S., Mehta, A. D., & Mesgarani, N. (2023). Joint, distributed and hierarchically organized encoding of linguistic features in the human auditory cortex. Nature Human Behaviour, 7(5), 740–753. 10.1038/s41562-023-01520-0

Kösem, A., Basirat, A., Azizi, L., & van Wassenhove, V. (2016). High-frequency neural activity predicts word parsing in ambiguous speech streams. Journal of Neurophysiology, 116(6), 2497–2512. 10.1152/jn.00074.2016

Kösem, A., Dai, B., McQueen, J. M., & Hagoort, P. (2023). Neural tracking of speech envelope does not unequivocally reflect intelligibility. NeuroImage, 272, 120040. 10.1016/j.neuroimage.2023.120040

Kösem, A., & van Wassenhove, V. (2017). Distinct contributions of low- and high-frequency neural oscillations to speech comprehension. Language, Cognition and Neuroscience, 32(5), 536–544. 10.1080/23273798.2016.1238495

Koutsoukos, E., Angelopoulos, E., Maillis, A., Papadimitriou, G. N., & Stefanis, C. (2013). Indication of increased phase coupling between theta and gamma EEG rhythms associated with the experience of auditory verbal hallucinations. Neuroscience Letters, 534, 242–245. 10.1016/j.neulet.2012.12.005

Kulasingham, J. P., Brodbeck, C., Presacco, A., Kuchinsky, S. E., Anderson, S., & Simon, J. Z. (2020). High gamma cortical processing of continuous speech in younger and older listeners. NeuroImage, 222, 117291. 10.1016/j.neuroimage.2020.117291

Lakatos, P., Karmos, G., Mehta, A. D., Ulbert, I., & Schroeder, C. E. (2008). Entrainment of neuronal oscillations as a mechanism of attentional selection. Science (New York, N.Y.), 320(5872), 110–113. 10.1126/science.1154735

Lakatos, P., Shah, A. S., Knuth, K. H., Ulbert, I., Karmos, G., & Schroeder, C. E. (2005). An Oscillatory Hierarchy Controlling Neuronal Excitability and Stimulus Processing in the Auditory Cortex. Journal of Neurophysiology, 94(3), 1904–1911. 10.1152/jn.00263.2005

Lalor, E., & Nidiffer, A. (2025). On the generative mechanisms underlying the cortical tracking of natural speech: A position paper. OSF. 10.31219/osf.io/xf8ay_v1

Lehtelä, L., Salmelin, R., & Hari, R. (1997). Evidence for reactive magnetic 10-Hz rhythm in the human auditory cortex. Neuroscience Letters, 222(2), 111–114. 10.1016/S0304-3940(97)13361-4

Lerousseau, J. P., Trébuchon, A., Morillon, B., & Schön, D. (2021). Frequency Selectivity of Persistent Cortical Oscillatory Responses to Auditory Rhythmic Stimulation. Journal of Neuroscience, 41(38), 7991–8006. 10.1523/JNEUROSCI.0213-21.2021

Leszczyński, M., Barczak, A., Kajikawa, Y., Ulbert, I., Falchier, A. Y., Tal, I., Haegens, S., Melloni, L., Knight, R. T., & Schroeder, C. E. (2020). Dissociation of broadband high-frequency activity and neuronal firing in the neocortex. Science Advances, 6(33), eabb0977. 10.1126/sciadv.abb0977

Lisman, J. E., & Idiart, M. A. (1995). Storage of 7 +/- 2 short-term memories in oscillatory subcycles. Science (New York, N.Y.), 267(5203), 1512–1515. 10.1126/science.7878473

Lisman, J. E., & Jensen, O. (2013). The θ-γ neural code. Neuron, 77(6), 1002–1016. 10.1016/j.neuron.2013.03.007

Lizarazu, M., Carreiras, M., & Molinaro, N. (2023). Theta-gamma phase-amplitude coupling in auditory cortex is modulated by language proficiency. Human Brain Mapping, 44(7), 2862–2872. 10.1002/hbm.26250

Lizarazu, M., Lallier, M., & Molinaro, N. (2019). Phase-amplitude coupling between theta and gamma oscillations adapts to speech rate. Annals of the New York Academy of Sciences, 1453(1), 140–152. 10.1111/nyas.14099

López-Madrona, V. J., Pérez-Montoyo, E., Álvarez-Salvado, E., Moratal, D., Herreras, O., Pereda, E., Mirasso, C. R., & Canals, S. (2020). Different theta frameworks coexist in the rat hippocampus and are coordinated during memory-guided and novelty tasks. eLife, 9, e57313. 10.7554/eLife.57313

López-Madrona, V. J., Trébuchon, A., Bénar, C. G., Schön, D., & Morillon, B. (2024). Different sustained and induced alpha oscillations emerge in the human auditory cortex during sound processing. Communications Biology, 7(1), 1–13. 10.1038/s42003-024-07297-w

López-Madrona, V. J., Trébuchon, A., Mindruta, I., Barbeau, E. J., Barborica, A., Pistol, C., Oane, I., Alario, F. X., & Bénar, C. G. (2024). Identification of early hippocampal dynamics during recognition memory with independent component analysis. eNeuro, 11(4). 10.1523/ENEURO.0183-23.2023

Lubinus, C., Keitel, A., Obleser, J., Poeppel, D., & Rimmele, J. M. (2023). Explaining flexible continuous speech comprehension from individual motor rhythms. Proceedings of the Royal Society B: Biological Sciences, 290(1994), 20222410. 10.1098/rspb.2022.2410

Lubinus, C., Orpella, J., Keitel, A., Gudi-Mindermann, H., Engel, A. K., Roeder, B., & Rimmele, J. M. (2021). Data-Driven Classification of Spectral Profiles Reveals Brain Region-Specific Plasticity in Blindness. Cerebral Cortex, 31(5), 2505–2522. 10.1093/cercor/bhaa370

Luo, H., & Poeppel, D. (2007). Phase Patterns of Neuronal Responses Reliably Discriminate Speech in Human Auditory Cortex. Neuron, 54(6), 1001–1010. 10.1016/j.neuron.2007.06.004

MacNeilage, P. F. (1998). The frame/content theory of evolution of speech production. Behavioral and Brain Sciences, 21(4), 499–511. 10.1017/S0140525X98001265

Mahjoory, K., Schoffelen, J.-M., Keitel, A., & Gross, J. (2020). The frequency gradient of human resting-state brain oscillations follows cortical hierarchies. eLife, 9, e53715. 10.7554/eLife.53715

Mai, G., Minett, J. W., & Wang, W. S.-Y. (2016). Delta, theta, beta, and gamma brain oscillations index levels of auditory sentence processing. NeuroImage, 133, 516–528. 10.1016/j.neuroimage.2016.02.064

Mäkelä, J. P., & Hari, R. (1987). Evidence for cortical origin of the 40 Hz auditory evoked response in man. Electroencephalography and Clinical Neurophysiology, 66(6), 539–546. 10.1016/0013-4694(87)90101-5

Marchesotti, S., Nicolle, J., Merlet, I., Arnal, L. H., Donoghue, J. P., & Giraud, A.-L. (2020). Selective enhancement of low-gamma activity by tACS improves phonemic processing and reading accuracy in dyslexia. PLoS Biology, 18(9), e3000833. 10.1371/journal.pbio.3000833

Mellem, M. S., Wohltjen, S., Gotts, S. J., Ghuman, A. S., & Martin, A. (2017). Intrinsic frequency biases and profiles across human cortex. Journal of Neurophysiology, 118(5), 2853–2864. 10.1152/jn.00061.2017

Mercier, M. R., Dubarry, A.-S., Tadel, F., Avanzini, P., Axmacher, N., Cellier, D., Vecchio, M. D., Hamilton, L. S., Hermes, D., Kahana, M. J., Knight, R. T., Llorens, A., Megevand, P., Melloni, L., Miller, K. J., Piai, V., Puce, A., Ramsey, N. F., Schwiedrzik, C. M., … Oostenveld, R. (2022). Advances in human intracranial electroencephalography research, guidelines and good practices. NeuroImage, 260, 119438. 10.1016/j.neuroimage.2022.119438

Mesgarani, N., Cheung, C., Johnson, K., & Chang, E. F. (2014). Phonetic Feature Encoding in Human Superior Temporal Gyrus. Science, 343(6174), 1006–1010. 10.1126/science.1245994

Meyer, L., Sun, Y., & Martin, A. E. (2020). Synchronous, but not entrained: Exogenous and endogenous cortical rhythms of speech and language processing. Language, Cognition and Neuroscience, 35(9), 1089–1099. 10.1080/23273798.2019.1693050

Michelmann, S., Treder, M. S., Griffiths, B., Kerrén, C., Roux, F., Wimber, M., Rollings, D., Sawlani, V., Chelvarajah, R., Gollwitzer, S., Kreiselmeyer, G., Hamer, H., Bowman, H., Staresina, B., & Hanslmayr, S. (2018). Data-driven re-referencing of intracranial EEG based on independent component analysis (ICA). Journal of Neuroscience Methods, 307, 125–137. 10.1016/j.jneumeth.2018.06.021

Molinaro, N., & Lizarazu, M. (2018). Delta(but not theta)-band cortical entrainment involves speech-specific processing. European Journal of Neuroscience, 48(7), 2642–2650. 10.1111/ejn.13811

Morillon, B., Lehongre, K., Frackowiak, R. S. J., Ducorps, A., Kleinschmidt, A., Poeppel, D., & Giraud, A.-L. (2010). Neurophysiological origin of human brain asymmetry for speech and language. Proceedings of the National Academy of Sciences of the United States of America, 107(43), 18688–18693. 10.1073/pnas.1007189107

Morillon, B., Liégeois-Chauvel, C., Arnal, L. H., Bénar, C.-G., & Giraud, A.-L. (2012). Asymmetric Function of Theta and Gamma Activity in Syllable Processing: An Intra-Cortical Study. Frontiers in Psychology, 3. 10.3389/fpsyg.2012.00248

Mukamel, R., Gelbard, H., Arieli, A., Hasson, U., Fried, I., & Malach, R. (2005). Coupling Between Neuronal Firing, Field Potentials, and fMRI in Human Auditory Cortex. Science, 309(5736), 951–954. 10.1126/science.1110913

Neymotin, S. A., Tal, I., Barczak, A., O’Connell, M. N., McGinnis, T., Markowitz, N., Espinal, E., Griffith, E., Anwar, H., Dura-Bernal, S., Schroeder, C. E., Lytton, W. W., Jones, S. R., Bickel, S., & Lakatos, P. (2022). Detecting Spontaneous Neural Oscillation Events in Primate Auditory Cortex. eNeuro, 9(4). 10.1523/ENEURO.0281-21.2022

Niedermeyer, E. (1990). Alpha-Like Rhythmical Activity of the Temporal Lobe. Clinical Electroencephalography, 21(4), 210–224. 10.1177/155005949002100410

Norman-Haignere, S. V., Keshishian, M., Devinsky, O., Doyle, W., McKhann, G. M., Schevon, C. A., Flinker, A., & Mesgarani, N. (2025). Temporal integration in human auditory cortex is predominantly yoked to absolute time. Nature Neuroscience, 28(11), 2356–2365. 10.1038/s41593-025-02060-8

Nourski, K. V. (2017). Auditory processing in the human cortex: An intracranial electrophysiology perspective. Laryngoscope Investigative Otolaryngology, 2(4), 147–156. 10.1002/lio2.73

Nourski, K. V., & Brugge, J. F. (2011). Representation of temporal sound features in the human auditory cortex. 22(2), 187–203. 10.1515/rns.2011.016

Nourski, K. V., Steinschneider, M., Rhone, A. E., Oya, H., Kawasaki, H., Howard, M. A., & McMurray, B. (2015). Sound identification in human auditory cortex: Differential contribution of local field potentials and high gamma power as revealed by direct intracranial recordings. Brain and language, 148, 37–50. 10.1016/j.bandl.2015.03.003

O’Dell, M. L., & Nieminen, T. (1999). Coupled oscillator model of speech rhythm. Proceedings of the XIVth International Congress of Phonetic Sciences, 2, 1075–1078.

Oganian, Y., & Chang, E. F. (2019). A speech envelope landmark for syllable encoding in human superior temporal gyrus. Science Advances, 5(11), eaay6279. 10.1126/sciadv.aay6279

Oganian, Y., Kojima, K., Breska, A., Cai, C., Findlay, A., Chang, E. F., & Nagarajan, S. S. (2023). Phase Alignment of Low-Frequency Neural Activity to the Amplitude Envelope of Speech Reflects Evoked Responses to Acoustic Edges, Not Oscillatory Entrainment. Journal of Neuroscience, 43(21), 3909–3921. 10.1523/JNEUROSCI.1663-22.2023

Park, H., Ince, R. A. A., Schyns, P. G., Thut, G., & Gross, J. (2015). Frontal Top-Down Signals Increase Coupling of Auditory Low-Frequency Oscillations to Continuous Speech in Human Listeners. Current Biology, 25(12), 1649–1653. 10.1016/j.cub.2015.04.049

Peelle, J. E., Gross, J., & Davis, M. H. (2013). Phase-Locked Responses to Speech in Human Auditory Cortex are Enhanced During Comprehension. Cerebral Cortex, 23(6), 1378–1387. 10.1093/cercor/bhs118

Pefkou, M., Arnal, L. H., Fontolan, L., & Giraud, A.-L. (2017). θ-Band and β-Band Neural Activity Reflects Independent Syllable Tracking and Comprehension of Time-Compressed Speech. Journal of Neuroscience, 37(33), 7930–7938. 10.1523/JNEUROSCI.2882-16.2017

Pellegrino, F., Coupé, C., & Marsico, E. (2011). A Cross-Language Perspective on Speech Information Rate. Language, 87(3), 539–558.

Picton, T. W., John, M. S., Dimitrijevic, A., & Purcell, D. (2003). Human auditory steady-state responses: Respuestas auditivas de estado estable en humanos. International Journal of Audiology, 42(4), 177–219. 10.3109/14992020309101316

Poeppel, D. (2001). Pure word deafness and the bilateral processing of the speech code. Cognitive Science, 25(5), 679–693. 10.1016/S0364-0213(01)00050-7

Poeppel, D. (2003). The analysis of speech in different temporal integration windows: Cerebral lateralization as ‘asymmetric sampling in time’. Speech Communication, 41(1), 245–255. 10.1016/S0167-6393(02)00107-3

Poeppel, D., & Assaneo, M. F. (2020). Speech rhythms and their neural foundations. Nature Reviews Neuroscience, 21(6), 322–334. 10.1038/s41583-020-0304-4

Pöppel, E. (1997). A hierarchical model of temporal perception. Trends in Cognitive Sciences, 1(2), 56–61. 10.1016/S1364-6613(97)01008-5

Quyen, M. L. V., Staba, R., Bragin, A., Dickson, C., Valderrama, M., Fried, I., & Engel, J. (2010). Large-Scale Microelectrode Recordings of High-Frequency Gamma Oscillations in Human Cortex during Sleep. Journal of Neuroscience, 30(23), 7770–7782. 10.1523/JNEUROSCI.5049-09.2010

Ramkumar, P., Parkkonen, L., & Hyvärinen, A. (2014). Group-level spatial independent component analysis of Fourier envelopes of resting-state MEG data. NeuroImage, 86, 480–491. 10.1016/j.neuroimage.2013.10.032

Rimmele, J. M., Poeppel, D., & Ghitza, O. (2021). Acoustically Driven Cortical δ Oscillations Underpin Prosodic Chunking. eNeuro, 8(4). 10.1523/ENEURO.0562-20.2021

Rosen, S. (1992). Temporal information in speech: Acoustic, auditory and linguistic aspects. *Philosophical Transactions of the Royal Society of London Series B*, Biological Sciences, 336(1278), 367–373. 10.1098/rstb.1992.0070

Schmidt, F., Chen, Y.-P., Keitel, A., Rösch, S., Hannemann, R., Serman, M., Hauswald, A., & Weisz, N. (2023). Neural speech tracking shifts from the syllabic to the modulation rate of speech as intelligibility decreases. Psychophysiology, 60(11), e14362. 10.1111/psyp.14362

Shannon, R. V., Zeng, F.-G., Kamath, V., Wygonski, J., & Ekelid, M. (1995). Speech Recognition with Primarily Temporal Cues. Science, 270(5234), 303–304. 10.1126/science.270.5234.303

Steinschneider, M., Nourski, K. V., & Fishman, Y. I. (2013). Representation of speech in human auditory cortex: Is it special? Hearing Research, 305, 57–73. 10.1016/j.heares.2013.05.013

Steriade, M. (2006). Grouping of brain rhythms in corticothalamic systems. Neuroscience, 137(4), 1087–1106. 10.1016/j.neuroscience.2005.10.029

Teng, X., & Poeppel, D. (2020). Theta and Gamma Bands Encode Acoustic Dynamics over Wide-Ranging Timescales. Cerebral Cortex (New York, N.Y.: 1991), 30(4), 2600–2614. 10.1093/cercor/bhz263

Teng, X., Tian, X., & Poeppel, D. (2016). Testing multi-scale processing in the auditory system. Scientific Reports, 6(1), 34390. 10.1038/srep34390

Teng, X., Tian, X., Rowland, J., & Poeppel, D. (2017). Concurrent temporal channels for auditory processing: Oscillatory neural entrainment reveals segregation of function at different scales. PLOS Biology, 15(11), e2000812. 10.1371/journal.pbio.2000812

Tiihonen, J., Hari, R., Kajola, M., Karhu, J., Ahlfors, S., & Tissari, S. (1991). Magnetoencephalographic 10-Hz rhythm from the human auditory cortex. Neuroscience Letters, 129(2), 303–305. 10.1016/0304-3940(91)90486-D

Tilsen, S. (2016). Selection and coordination: The articulatory basis for the emergence of phonological structure. Journal of Phonetics, 55, 53–77. 10.1016/j.wocn.2015.11.005

Vanthornhout, J., Decruy, L., Wouters, J., Simon, J. Z., & Francart, T. (2018). Speech Intelligibility Predicted from Neural Entrainment of the Speech Envelope. Journal of the Association for Research in Otolaryngology: JARO, 19(2), 181–191. 10.1007/s10162-018-0654-z

Varnet, L., Ortiz-Barajas, M. C., Erra, R. G., Gervain, J., & Lorenzi, C. (2017). A cross-linguistic study of speech modulation spectra. The Journal of the Acoustical Society of America, 142(4), 1976–1989. 10.1121/1.5006179

Wang, J., Gao, D., Li, D., Desroches, A. S., Liu, L., & Li, X. (2014). Theta–gamma coupling reflects the interaction of bottom-up and top-down processes in speech perception in children. NeuroImage, 102, 637–645. 10.1016/j.neuroimage.2014.08.030

Wang, X., Delgado, J., Marchesotti, S., Kojovic, N., Sperdin, H. F., Rihs, T. A., Schaer, M., & Giraud, A.-L. (2023). Speech Reception in Young Children with Autism Is Selectively Indexed by a Neural Oscillation Coupling Anomaly. The Journal of Neuroscience: The Official Journal of the Society for Neuroscience, 43(40), 6779–6795. 10.1523/JNEUROSCI.0112-22.2023

Wang, Y., Wu, D., Ding, N., Zou, J., Lu, Y., Ma, Y., Zhang, X., Yu, W., & Wang, K. (2025). Linear phase property of speech envelope tracking response in Heschl’s gyrus and superior temporal gyrus. Cortex, 186, 1–10. 10.1016/j.cortex.2025.02.015

Xu, N., Zhao, B., Luo, L., Zhang, K., Shao, X., Luan, G., Wang, Q., Hu, W., & Wang, Q. (2023). Two stages of speech envelope tracking in human auditory cortex modulated by speech intelligibility. Cerebral Cortex, 33(5), 2215–2228. 10.1093/cercor/bhac203

## Supplementary bibliography

Arinyo-i-Prats, A., López-Madrona, V. J., & Paluš, M. (2024). Lead/Lag directionality is not generally equivalent to causality in nonlinear systems: Comparison of phase slope index and conditional mutual information. NeuroImage, 120610. 10.1016/j.neuroimage.2024.120610

Artoni, F., Delorme, A., & Makeig, S. (2018). Applying dimension reduction to EEG data by Principal Component Analysis reduces the quality of its subsequent Independent Component decomposition. NeuroImage, 175, 176–187. 10.1016/j.neuroimage.2018.03.016

Aru, J., Aru, J., Priesemann, V., Wibral, M., Lana, L., Pipa, G., Singer, W., & Vicente, R. (2015). Untangling cross-frequency coupling in neuroscience. Current Opinion in Neurobiology, 31, 51–61. 10.1016/j.conb.2014.08.002

Barnett, L., & Seth, A. K. (2014). The MVGC multivariate Granger causality toolbox: A new approach to Granger-causal inference. Journal of Neuroscience Methods, 223, 50–68. 10.1016/j.jneumeth.2013.10.018

Bell, A. J., & Sejnowski, T. J. (1995). An information-maximization approach to blind separation and blind deconvolution. Neural Computation, 7(6), 1129–1159. 10.1162/neco.1995.7.6.1129

Bressler, S. L., & Seth, A. K. (2011). Wiener-Granger causality: A well established methodology. NeuroImage, 58(2), 323–329. 10.1016/j.neuroimage.2010.02.059

Canolty, R. T., Edwards, E., Dalal, S. S., Soltani, M., Nagarajan, S. S., Kirsch, H. E., Berger, M. S., Barbaro, N. M., & Knight, R. T. (2006). High Gamma Power Is Phase-Locked to Theta Oscillations in Human Neocortex. Science, 313(5793), 1626–1628. 10.1126/science.1128115

Cohen, M. X. (2014). Analyzing Neural Time Series Data: Theory and Practice. MIT Press.

Cole, S., & Voytek, B. (2019). Cycle-by-cycle analysis of neural oscillations. Journal of Neurophysiology. 10.1152/jn.00273.2019

Delorme, A., & Makeig, S. (2004). EEGLAB: An open source toolbox for analysis of single-trial EEG dynamics including independent component analysis. Journal of Neuroscience Methods, 134(1), 9–21. 10.1016/j.jneumeth.2003.10.009

Dupré la Tour, T., Tallot, L., Grabot, L., Doyère, V., van Wassenhove, V., Grenier, Y., & Gramfort, A. (2017). Non-linear auto-regressive models for cross-frequency coupling in neural time series. PLoS Computational Biology, 13(12), e1005893. 10.1371/journal.pcbi.1005893

Geweke, J. (1982). Measurement of Linear Dependence and Feedback between Multiple Time Series. Journal of the American Statistical Association, 77(378), 304–313. 10.1080/01621459.1982.10477803

Gilbert, G., & Lorenzi, C. (2006). The ability of listeners to use recovered envelope cues from speech fine structure. The Journal of the Acoustical Society of America, 119(4), 2438–2444. 10.1121/1.2173522

Granger, C. W. J. (1969). Investigating Causal Relations by Econometric Models and Cross-spectral Methods. Econometrica, 37(3), 424–438. 10.2307/1912791

Groppe, D. M., Bickel, S., Dykstra, A. R., Wang, X., Mégevand, P., Mercier, M. R., Lado, F. A., Mehta, A. D., & Honey, C. J. (2017). iELVis: An open source MATLAB toolbox for localizing and visualizing human intracranial electrode data. Journal of Neuroscience Methods, 281, 40–48. 10.1016/j.jneumeth.2017.01.022

GTZAN Dataset—Music Genre Classification. (s. f.). Recuperado 16 de mayo de 2025, de https://www.kaggle.com/datasets/andradaolteanu/gtzan-dataset-music-genre-classification

Herreras, O., Torres, D., Martín-Vázquez, G., Hernández-Recio, S., López-Madrona, V. J., Benito, N., Makarov, V. A., & Makarova, J. (2022). Site-dependent shaping of field potential waveforms. Cerebral Cortex, bhac297. 10.1093/cercor/bhac297

Kisler, T., Reichel, U., & Schiel, F. (2017). Multilingual processing of speech via web services. Computer Speech & Language, 45, 326–347. 10.1016/j.csl.2017.01.005

Kroos, C., Bones, O., Cao, Y., Harris, L., Jackson, P., Davies, W., Wang, W., Cox, T., & Plumbley, M. (2019). Generalisation in environmental sound classification: The ‘making sense of sounds’ data set and challenge. ICASSP 2019-2019 IEEE International Conference on Acoustics, Speech and Signal Processing (ICASSP).

López-Madrona, V. J., Matias, F. S., Mirasso, C. R., Canals, S., & Pereda, E. (2019). Inferring correlations associated to causal interactions in brain signals using autoregressive models. Scientific Reports, 9(1), 17041. 10.1038/s41598-019-53453-2

Makarova, J., Toledano, R., Blázquez-Llorca, L., Sánchez-Herráez, E., Gil-Nagel, A., DeFelipe, J., & Herreras, O. (2025). Intracranial Voltage Profiles from Untangled Human Deep Sources Reveal Multisource Composition and Source Allocation Bias. Journal of Neuroscience, 45(1). 10.1523/JNEUROSCI.0695-24.2024

Medina Villalon, S., Paz, R., Roehri, N., Lagarde, S., Pizzo, F., Colombet, B., Bartolomei, F., Carron, R., & Bénar, C.-G. (2018). EpiTools, A software suite for presurgical brain mapping in epilepsy: Intracerebral EEG. Journal of Neuroscience Methods, 303, 7–15. 10.1016/j.jneumeth.2018.03.018

Nolte, G., Ziehe, A., Krämer, N., Popescu, F., & Müller, K.-R. (2010). Comparison of Granger Causality and Phase Slope Index. Causality: Objectives and Assessment, 267–276. http://proceedings.mlr.press/v6/nolte10a.html

Nolte, G., Ziehe, A., Nikulin, V. V., Schlögl, A., Krämer, N., Brismar, T., & Müller, K.-R. (2008). Robustly estimating the flow direction of information in complex physical systems. Physical Review Letters, 100(23), 234101. 10.1103/PhysRevLett.100.234101

Oostenveld, R., Fries, P., Maris, E., & Schoffelen, J.-M. (2011). FieldTrip: Open source software for advanced analysis of MEG, EEG, and invasive electrophysiological data. Computational Intelligence and Neuroscience, 2011, 156869. 10.1155/2011/156869

Penny, W., Friston, K., Ashburner, J., Kiebel, S., & Nichols, T. (2007). Statistical Parametric Mapping: The Analysis of Functional Brain Images. 10.1016/B978-0-12-372560-8.X5000-1

Podvalny, E., Noy, N., Harel, M., Bickel, S., Chechik, G., Schroeder, C. E., Mehta, A. D., Tsodyks, M., & Malach, R. (2015). A unifying principle underlying the extracellular field potential spectral responses in the human cortex. Journal of Neurophysiology, 114(1), 505–519. 10.1152/jn.00943.2014

Radford, A., Kim, J. W., Xu, T., Brockman, G., McLeavey, C., & Sutskever, I. (2022). Robust Speech Recognition via Large-Scale Weak Supervision (arXiv:2212.04356). arXiv. 10.48550/arXiv.2212.04356

Stolk, A., Griffin, S., van der Meij, R., Dewar, C., Saez, I., Lin, J. J., Piantoni, G., Schoffelen, J.-M., Knight, R. T., & Oostenveld, R. (2018). Integrated analysis of anatomical and electrophysiological human intracranial data. Nature Protocols, 13(7), 1699–1723. 10.1038/s41596-018-0009-6

Talairach, J., Tournoux, P., Musolino, A., & Missir, O. (1992). Stereotaxic exploration in frontal epilepsy. Advances in Neurology, 57, 651–688.

Thomson, D. J. (1982). Spectrum estimation and harmonic analysis. Proceedings of the IEEE, 70(9), 1055–1096. Proceedings of the IEEE. 10.1109/PROC.1982.12433

Tort, A. B. L., Kramer, M. A., Thorn, C., Gibson, D. J., Kubota, Y., Graybiel, A. M., & Kopell, N. J. (2008). Dynamic cross-frequency couplings of local field potential oscillations in rat striatum and hippocampus during performance of a T-maze task. Proceedings of the National Academy of Sciences of the United States of America, 105(51), 20517–20522. 10.1073/pnas.0810524105

Voytek, B., Kramer, M. A., Case, J., Lepage, K. Q., Tempesta, Z. R., Knight, R. T., & Gazzaley, A. (2015). Age-Related Changes in 1/f Neural Electrophysiological Noise. Journal of Neuroscience, 35(38), 13257–13265. 10.1523/JNEUROSCI.2332-14.2015

Zanon Boito, M., Havard, W., Garnerin, M., Le Ferrand, É., & Besacier, L. (2020). MaSS: A Large and Clean Multilingual Corpus of Sentence-aligned Spoken Utterances Extracted from the Bible. En N. Calzolari, F. Béchet, P. Blache, K. Choukri, C. Cieri, T. Declerck, S. Goggi, H. Isahara, B. Maegaard, J. Mariani, H. Mazo, A. Moreno, J. Odijk, & S. Piperidis (Eds.), Proceedings of the Twelfth Language Resources and Evaluation Conference (pp. 6486–6493). European Language Resources Association. https://aclanthology.org/2020.lrec-1.799/

